# Complex slow waves radically reorganise human brain dynamics under 5-MeO-DMT

**DOI:** 10.1101/2024.10.04.616717

**Authors:** George Blackburne, Rosalind G. McAlpine, Marco Fabus, Alberto Liardi, Sunjeev K. Kamboj, Pedro A. M. Mediano, Jeremy I. Skipper

## Abstract

5-methoxy-N,N-dimethyltryptamine (5-MeO-DMT) is a psychedelic drug known for its uniquely profound effects on subjective experience, reliably eradicating the perception of time, space, and the self. However, little is known about how this drug alters large-scale brain activity. We collected naturalistic electroencephalography (EEG) data of 29 healthy individuals before and after inhaling a high dose (12mg) of vaporised synthetic 5-MeO-DMT. We replicate work from rodents showing amplified low-frequency oscillations, but extend these findings with novel tools for characterising the organisation and dynamics of complex low-frequency spatiotemporal fields of neural activity. We find that 5-MeO-DMT radically reorganises low-frequency flows of neural activity, causing them to become incoherent, heterogeneous, viscous, fleeting, nonrecurring, and to cease their typical travelling forwards and backwards across the cortex compared to resting state. Further, we find a consequence of this reorganisation in broadband activity, which exhibits slower, more stable, low-dimensional behaviour, with increased energy barriers to rapid global shifts. These findings provide the first detailed empirical account of how 5-MeO-DMT sculpts human brain dynamics, revealing a novel set of cortical slow wave behaviours, with significant implications for extant neuroscientific models of serotonergic psychedelics.

## 1 Introduction

Psychedelic drugs are reentering scientific research as unique and reliable tools for interrogating the neural processes that entail alterations in the structure of subjective experience [1, 2]. In tandem, they are being explored as potential treatments for an array of mental health conditions [3, 4]. However, 5-methoxy-N,N-dimethyltryptamine (5-MeO-DMT), likely the most profound agent in this class [5], has remained understudied in human neuroscience. This is despite its current involvement in clinical trials for the treatment of depression [6, 7], bipolar [8], and alcohol use disorders [9].

5-MeO-DMT is a naturally occurring, potent, rapid-acting tryptamine, with use likely dating back over 1000 years in South American Tiwanaku ritual practices via insufflation of the trace concentrations contained in Anadenanthera seeds [10, 11]. Contemporary usage however, is markedly different, predominantly occurring through either the synthetic form, or via the venom secreted from the parotid glands of the Incilius alvarius toad [12]. Indeed, it is the high-dose combustion or vaporisation and inhalation of these formats through which 5-MeO-DMT has attained its unofficial status as the world’s most powerful psychedelic, or the ‘God Molecule’ [5]. The experience elicited by inhaled 5-MeO-DMT, like its chemical relative DMT, is characterised by a radical progression, or ‘breakthrough’, into what is felt as ‘another dimension’, such that sensation becomes detached from consensus reality [13]. However, unlike DMT, this environmental disconnection generally does not typically involve immersion into an overwhelmingly visually intricate geometric space filled with seemingly sentient entities, but rather a world entirely lacking the usual axes of variation to structure it. At the peak of the 5-MeO-DMT experience, rudimentary structures of subjective experience such as time, space and the self are felt to be abolished, and a dramatic lack of differentiation ensues [14]. Fully awake and aware, individuals feel they have entered into an ineffable realm of ‘everything and nothing’, where even the inference that they are the subject of the experience is protracted [15]. 5-MeO-DMT is therefore a perturbation that renders which of a creature’s consciousness-related capacities can be online, but does not in the classical sense ‘reduce’ conscious level [16]. Thus, it may be most productively thought of as a *global mode of deconstructed consciousness*.

Preclinical work in awake rodents indicates that 5-MeO-DMT elicits its most pronounced effects in augmenting low-frequency rhythms (<4Hz) [17–20]. 5-MeO-DMT has been shown to increase the power of low-frequency oscillations in local field potentials, with concurrent multi-unit recordings showing neural activity to alternate between periods of high-firing and generalised silence, while mice freely behave. This hybrid state was therefore dubbed ‘paradoxical wakefulness’, since mice exhibited the canonical neurophysiological signatures of slow-wave sleep (SWS) while awake [19]. The amplification of low-frequency oscillations is thought to be a domain-general indicator of loss of consciousness beyond SWS, such as the period of slow-wave activity saturation (SWAS) in general anaesthesia [21], and coma patients being diagnosed with unresponsive wakefulness syndrome [22, 23]. Thus, these results with 5-MeO-DMT challenge the idea that amplified low-frequency rhythms have a one-to-one correspondence with reductions in conscious level [24].

Simple oscillatory phenomena in the brain are increasingly being empirically understood as univariate symptoms of more complex spatiotemporal propagating patterns, with SWS and SWAS exemplifying this shift in understanding [25–28]. Indeed, there have been recent theoretical and empirical calls to define brain states more generally via wave-like processes [29–31], though the idea that the brain is a non-stationary system composed of trajectories inextricably structured in space and time is not itself a recent development [32–34]. Complex flows, or metastable waves, are transient activity patterns that sweep across the cortex and converge around points of stationarity, termed singularities. These can take on a host of forms, such as sources or sinks, which expand or contract around a point, spirals which rotate around a point, and saddles which are usually the superposition of different interacting waves. These patterns have been consistently observed in mesoscopic and macroscopic neural recordings [30, 35–39], and are thought to play crucial roles in cortical computation [40, 41], modulating excitation-inhibition balance [42], memory [43, 44], and perception [45–47].

Thus, a productive approach for the empirical characterisation of the effects of 5-MeO-DMT on macroscopic brain dynamics might lie in understanding how complex low-frequency flows are reorganised. We hypothesised that, while 5-MeO-DMT may look analogous to states of unconsciousness from a univariate oscillatory perspective, it may look strikingly different from a more comprehensive spatiotemporal standpoint. Instead of inducing more coherent, durable and stereotyped global low-frequency flow structures, like in anaesthesia [25, 27, 48], we predicted that 5-MeO-DMT would induce a state of resolute complex low-frequency incoherence, with disjointed and viscous flows limiting the emergence of global propagating patterns able to spread across the cortex. We hypothesised that there would be a greater occurrence of localised low-frequency patterns with differing directions and speeds, possessing short lifetimes, and bearing little resemblance to each other over time. In sum, we expected that 5-MeO-DMT would upset the backbone of cortical dynamics with ‘broken’, rather than tidal, waves.

Importantly, one should ask what such low-frequency changes functionally imply for statistical motifs of broadband (wide frequency) brain dynamics. Owing to the expected spectral alterations, we predicted that the intrinsic timescales of the cortex would shift towards a slower regime, and that excursions from similar activity states would depart from each other more tardily. Crucially, if slow spatiotemporal flows are disorganised, the brain may be less able to effectively orchestrate large rapid global amplitude reconfigurations, thus increasing the steady-state residence and stability of broadband cortical dynamics [49]. Given this, broadband responses may be forced from their large population space onto an unusual and simpler sub-space. Put simply, these flows may curb the degrees of freedom of the brain. This bounding of the system’s dimensionality could distort and ablate the typical construction of neural representations [50], as such, the individual’s model of the world should collapse in parameters, resulting in the deconstruction of subjective experience.

We investigate these predictions by collecting naturalistic electroencephalography (EEG) data of individuals (N=29) before and immediately after inhaling a high-dose (12mg) of vaporised synthetic 5-MeO-DMT. We first replicate recent work in rodents on the power of low-frequency oscillations, but extend this by performing a suite of analyses characterising alterations in the organisation and dynamics of complex low-frequency spatiotemporal flows of neural activity. We then complement this by assessing the statistical behaviour of broadband signals across the cortex in terms of their intrinsic timescales, separation rates, underlying energy landscape, and dimensionality. In doing so, we provide the first detailed account of the neuroscience of 5-MeO-DMT in humans.

## 2 Results

### Slow oscillations

Given the lack of 5-MeO-DMT research in humans, we begin by assessing the frequency content of our 64-channel EEG data. The signature effect of 5-MeO-DMT in mice has been the amplification of low-frequency oscillations (*<*4 Hz). A peculiar effect, since augmentation of low-frequency content has hitherto been considered a signature of loss of wakefulness and awareness. Moreover, rodent studies have shown no clear effect of 5-MeO-DMT on the power of alpha oscillations, the deflation of which is currently considered the neurophysiological signature of psychedelic drug effects in humans [51–64]. To enable robust time-frequency representation of cortical signals, we use Empirical Mode Decomposition (EMD). We do so since canonical methods such as the Fourier Transform (FT), can provide unreliable estimates when signals are noisy, non-sinusoidal, and non-stationary, as is routinely seen in in neural recordings [65–70]. As an illustration, a slightly non-linear high-delta (4 Hz) oscillation can be mistaken to contain a low-alpha (8 Hz) rhythm if the FT is applied due to harmonic effects (Supplementary Fig.1). Therefore, we use EMD to iteratively sift out the intrinsic basis functions of cortical signals without assuming they are sinusoidal or static. Here, we apply EMD and the Hilbert-Huang Transform (HHT) to the epoched data in the range of 0.5-50Hz (n=19, 10 exlcuded to movement artifacts), to maximise the number of viable participants. The non-linear power spectrum is taken as the sum across instantaneous frequencies over intrinsic mode functions (IMFs), normalised by their density.

5-MeO-DMT shifts the power spectra of cortical signals at its antipodes. Slow (<1.5Hz) and fast (>20Hz) oscillations rapidly rise in power within 20s after drug inhalation and decay after 8-10 minutes (Fig.1a,d), coincident with the expected rise and decay of drug effects. Alterations in spectra can be seen to track the expected duration of drug effects, with the first 4 minutes being marked by a particularly constant spectral profile (Fig.1b).

**Figure 1.**
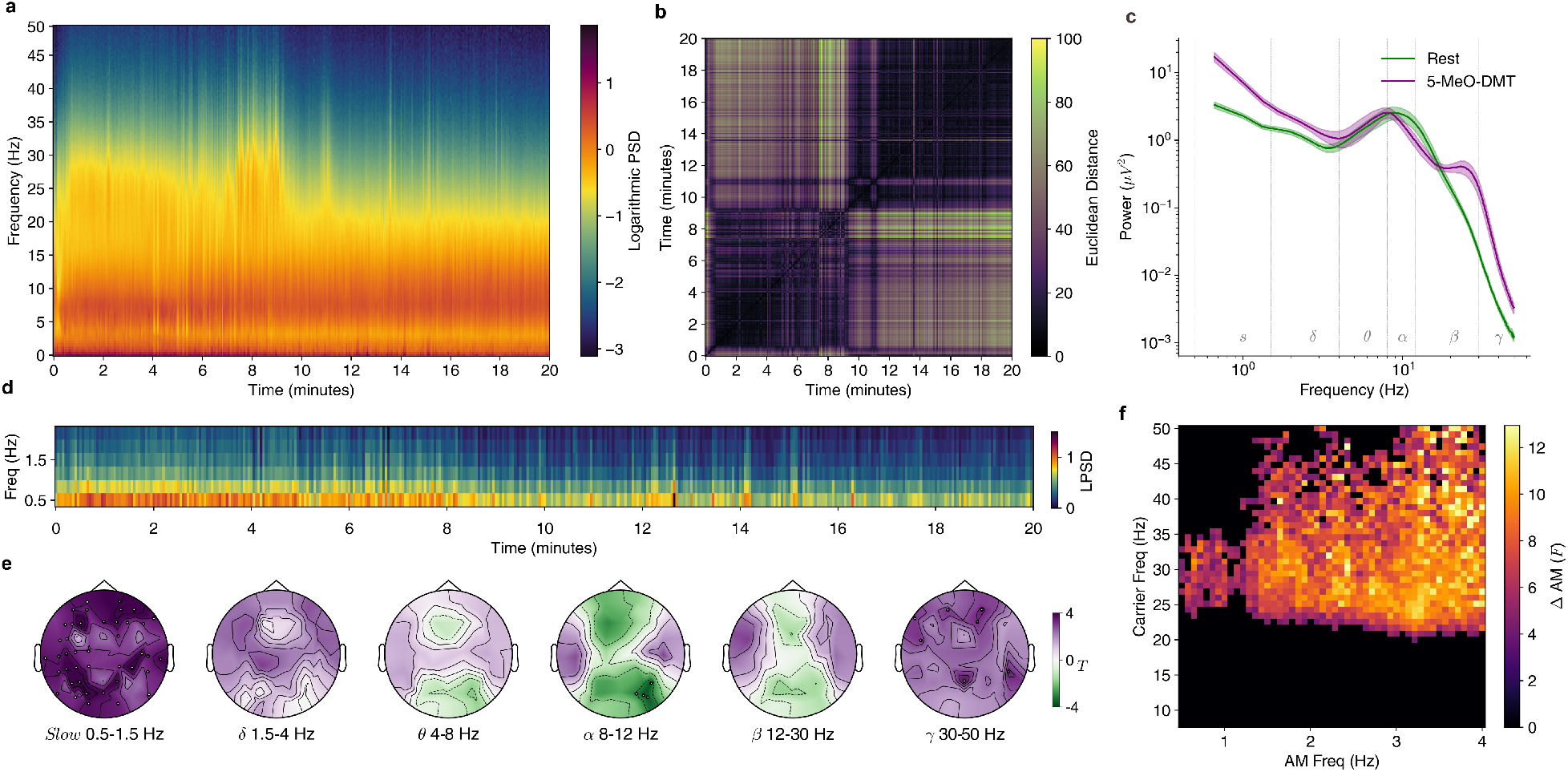
5-MeO-DMT induces diffuse high-amplitude slow oscillations. **a** Average time-frequency power spectral density (PSD) for the 20 minutes post-inhalation of 5-MeO-DMT, derived via Empirical Mode Decomposition and the Hilbert-Huang transform. **b** Euclidean distance matrix for average spectra vectors over time. **c** Time-averaged power spectra for eyes-closed baseline period and estimated peak 5-MeO-DMT at 1.5-2.5 minutes (mean *±* SEM). **d** Focused low-frequency spectrogram. **e** Scalp topographic statistical maps for T-test comparing the two conditions per frequency band and electrode, with electrodes representing significant differences (p<.05 FDR-corrected Benjamini–Hochberg procedure). Purple indicates increases under 5-MeO-DMT, green indicates reductions. **f** Holospectrum statistical map for ANOVA comparing the two conditions, with significant clusters coloured (p<.05 cluster-corrected).

Examining the estimated peak of the drug effect (1.5-2.5 minutes) reveals a stark effect at low-frequencies. We find that the mean power of the lowest frequency bin (0.5Hz), increases by 415% compared to baseline (Fig.1c). These amplified slow oscillations (0.5-1.5Hz) are spatially distributed, with increases occurring over frontal, parietal, temporal, central and occipital electrodes (Fig.1e; max: PO3 *T* =5.104, *p*_*FDR*_ =0.011, *BF*_10_ =365.623, *d* =1.472).

Inspecting higher-frequency changes, we observe distributed but weaker increases in gamma oscillations at the estimated peak (Fig.1e max: Pz *T* =3.750, *p*_*FDR*_ =0.001, *BF*_10_ =26.542, *d* =1.025). As the expected end-point of the experience is arriving, between 7.5-9 minutes, there is a sudden jump in high-frequency oscillations (Fig.1a). Notably, we see that alpha oscillations are not robustly reduced across every area of the cortex with the large effect sizes seen with other psychedelic compounds [63] (Fig.1c). In fact, when parameterising the peaks of the average power spectra with Gaussians [71], we find no significant change in alpha (*p*_*FDR*_ =0.411), but a significant reduction in the peak frequency (*p*_*FDR*_ =0.023). Examining the topography of alpha changes we find significant reductions over the right parietal-occipital cortex (Pz: *p*_*FDR*_ =0.030), but also increases in alpha power over the temporal lobes approaching significance (T8: *p*_*FDR*_ =0.052).

Finally, we investigate whether fast oscillations can themselves be understood in terms of slow oscillations. Specifically, we test if there is greater amplitude modulation of high-frequency content by low-frequency content under 5-MeO-DMT, as evidenced by changes in their holospectrum. Indeed, we find that high-frequency rhythms exhibit greater fluctuations in power at low-frequencies under 5-MeO-DMT (Fig.1f; p=.005 ANOVA cluster-level permutation).

In sum, these results provide evidence for increases in the power of slow-oscillations in the human brain under 5-MeO-DMT, replicating results from rodents [17–19], as well as providing evidence that substantial whole-brain loss of alpha power may not be as essential for the 5-MeO-DMT experience, in contrast to other classical psychedelics.

### Complex slow waves

Moving past characterising the power of univariate rhythmic fluctuations in neural activity in the two conditions, we characterise multivariate macroscopic propagating patterns with particular temporal motifs. We do this by adopting the mathematics of flow velocity from fluid dynamics. We first perform EMD on the continuous signals from the peak window for those participants who have it artefact free (1.5-2.5 minutes post-ingestion, n=13) to maximise detection of the low-frequency cycles, obtaining six time-evolving oscillatory intrinsic mode functions (IMFs), for each electrode [72]. These modes have a mean instantaneous frequency (IF) of 16.398*±*1.104, 6.797*±*0.290, 2.270*±*0.093, 0.741*±*0.028, 0.319*±*0.007, 0.133*±*0.004 respectively (all in Hz). Since our interest is low-frequency modes, we only analyse the third, fourth and fifth modes, excluding the sixth as a residual artefactual drift, and refer to these as Delta, Slow and Ultra-Slow throughout. The mean IF of these modes do not significantly change between conditions (Fig.2a; Delta: *T* =3.012, *p*_*FDR*_ =0.065; Slow: *T* =0.462, *p*_*FDR*_ =1; Ultra-Slow: *T* =-0.112, *p*_*FDR*_ =1). After interpolation of the instantaneous amplitude and phase of the modes onto a uniform 32×32 scalp grid, mimicking the pixels from optical imaging voltage grids [48], we obtain the velocity field at each sample for each mode and condition (see Methods). As expected, we detect continuous and complex propagating spatiotemporal patterns in both conditions, but with markedly different structure and dynamics (Fig.2b-m).

**Figure 2.**
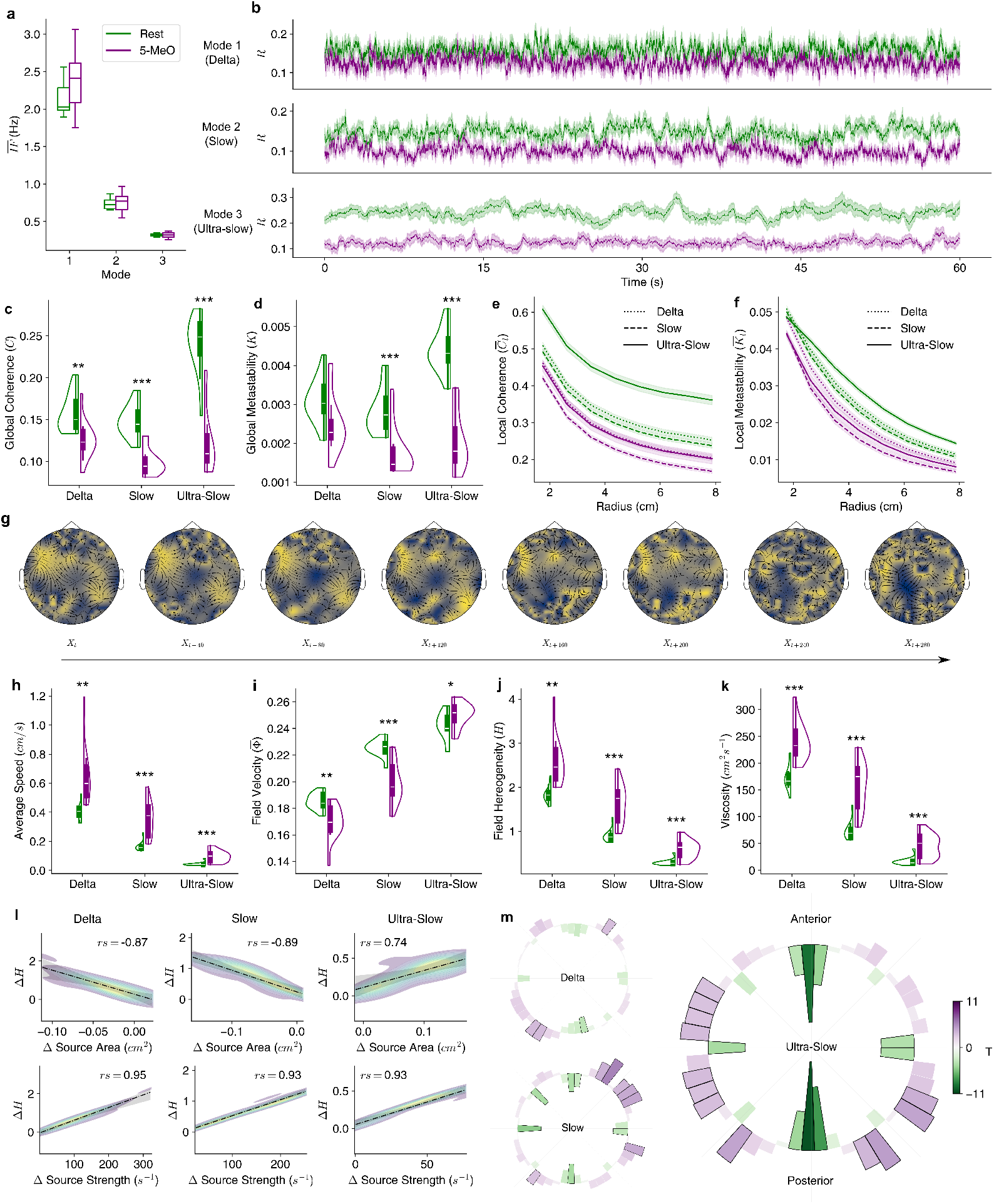
5-MeO-DMT reorganises patterns of complex low-frequency flow. **a** Mean instantaneous frequency for each intrinsic mode function (IMF). **b** Instantaneous global phase coherence for each IMF (mean *±* SEM). **c** Global phase coherence for each IMF. **d** Local phase coherence for each IMF across radii (mean *±* SEM). **e** Global phase metastability for each IMF. **f** Local phase metastability for each IMF across radii (mean *±* SEM). **g** Example velocity field for Mode 2 from a representative participant over the course of 560ms during estimated peak of 5-MeO-DMT. **h** Mean wave speed for each IMF. **i** Average normalised velocity of the velocity field for each IMF. **j** Average heterogeneity of the velocity field for each IMF. **k** Viscosity of the velocity field for each IMF. **l** Correlations between changes in field heterogeneity with changes in source area (top row) and source strength (bottom row) for each IMF (columns). Change scores are 5-MeO-DMT -Rest. **m** Changes in the direction of flow for each IMF across the cortex. Bin position indicates direction, and bin orientation indicates change. Inward pointing bins indicate reduced travel under 5-MeO-DMT. T-test p<.05 FDR-corrected bins are indicated with black borders. (^*^,^**^,^***^ indicates p<.05,.01,.001 FDR-corrected Benjamini–Hochberg procedure)

Specifically, assessing alterations in phase interactions of the field via a Kuramoto-like formalism (see Methods), we find that 5-MeO-DMT reduces the phase synchrony (global coherence) of low-frequency phase fields Fig.2c (Delta: *p*_*FDR*_ =0.010; Slow: *p*_*FDR*_ =2.07*×*10^−5^; Ultra-Slow: *p*_*FDR*_ =8.73*×*10^−7^), and also results in less variance in this synchrony over time (global metastability) (Fig.2d Delta: *p*_*FDR*_ =0.080; Slow: *p*_*FDR*_ =5.96*×*10^−4^; Ultra-Slow: *p*_*FDR*_ =8.73*×*10^−7^). We find that this is also the case locally, across radii ranging from 1.75-7.875 cm in increments of 0.875 cm for both coherence (Fig.2e), and metastability (Fig.2f) (see Supplementary Table.2 for statistics) Thus, 5-MeO-DMT induces a state of persistent low-frequency desynchronisation.

Examining the velocity fields further, we find that 5-MeO-DMT causes waves to travel faster on average (Fig.2h; Delta: *p*_*FDR*_ =0.002; Slow: *p*_*FDR*_ =1.49*×*10^−4^; Ultra-Slow: *p*_*FDR*_ =2.61*×*10^−4^). However, this largely reflects disorganised and spatially circumscribed flow, since globally the field exhibits less directional alignment and collective motion, as shown by reductions in the fields normalised velocity for Delta and Slow waves (Fig.2i; Delta: *p*_*FDR*_ =0.003; Slow: *p*_*FDR*_ =7.19*×*10^−5^). However, the opposite is true for Ultra-Slow waves (Fig.2i *p*_*FDR*_ =0.026). Supporting the idea that flow becomes more fragmented, we find that velocity fields become significantly more spatially heterogeneous (Fig.2j; Delta: *p*_*FDR*_ =0.001; Slow: *p*_*FDR*_ =1.49*×*10^−4^; Ultra-Slow: *p*_*FDR*_ =1.49*×*10^−4^). We predicted that this would be the result of decreases in the area occupied by flow sources, and increases in the strength (divergence) exerted by flow sources. Indeed, we show that such decreases in the area of sources and increases in the strength of sources strongly correlate with the observed changes in field heterogeneity, with the lowest *rs* = .74 (Fig.2l).

Finally, we further confirm the ‘provincialising’ effects of 5-MeO-DMT on low-frequency flow by showing it to become more viscous, revealing that the diffusion of neural activity across the cortex at this timescale is significantly hampered by the drug (Fig.2k; Delta: *p*_*FDR*_ =5.23*×*10^−4^; Slow: *p*_*FDR*_ =2.34*×*10^−4^; Ultra-Slow: *p*_*FDR*_ =7.35*×*10^−4^). In sum, 5-MeO-DMT induces a state of disorganised low-frequency flow, giving rise to greater local patterns with diverse shapes, directions and speeds. These flows collectively diminish the emergence of global propagating patterns with specific paths across the cortex.

To directly assess changes in the path of waves across the cortex, we obtain the angle of each grid element on the scalp via the inverse of the tangent of their X and Y directional vectors. We bin the 2*π* complete angle of directions into 50 bins, and find that 5-MeO-DMT significantly alters wave travel directionality (Fig.2m). We find that Slow and Ultra-Slow waves travel significantly less in true anterior, posterior directions, and lateral directions, instead preferring to travel in an intercardinal fashion (Fig.2m). The loss of canonical forward and backward travelling waves is particularly striking in the case of Ultra-Slow waves (Fig.2m; Anterior travel: *T* =-10.375, *p*_*FDR*_ =3.61*×*10^−5^, *BF*_10_ =6.11*×*10^4^, *d* =2.639; Posterior travel: *T* =-9.065, *p*_*FDR*_ =7.67*×*10^−5^, *BF*_10_ =1.66*×*10^4^, *d* =2.648). Thus, 5-MeO-DMT reorganises wave travel and specifically diminishes forwards and backward travelling slow waves.

Visual inspection of velocity fields at baseline indicated that field patterns were persistent and frequently reoccurred. However, under 5-MeO-DMT patterns appeared to have shorter dwell times, with future patterns noticeably more dissimilar to past ones. To specifically test such alterations in the temporal dynamics of low-frequency flows, we compute recurrence matrices for the velocity fields (see Methods). Fig.3a-b show example network plots for the first 1*/*4 (15s) of the recurrence network for Mode 2 from a representative participant. Fig.3c shows the same for the remaining participants. The 5-MeO-DMT recurrence networks can be seen to be scattered and less densely interconnected, as evidenced by the average degree distributions for each condition in (Fig.3d). The mean degree of the recurrence network is significantly reduced by 5-MeO-DMT for all three modes (Delta: *p*_*FDR*_ =1.38*×*10^−4^. Slow: *p*_*FDR*_ =4.21*×*10^−6^. Ultra-Slow: *p*_*FDR*_ =4.21*×*10^−6^). Indeed, we also see that the number of communities in the recurrence network significantly increases under 5-MeO-DMT (Fig.3e Delta: *p*_*FDR*_ =6.89*×*10^−5^; Slow: *p*_*FDR*_ =0.003; Ultra-Slow: *p*_*FDR*_ =0.033;), indicating an increase in the number of unique field patterns. Finally we see that the global efficiency of the network, representing how effectively flow patterns are exchanged between the past and the future, is significantly reduced by 5-MeO-DMT for all three modes (Delta: *p*_*FDR*_ =0.001. Slow: *p*_*FDR*_ =6.61*×*10^−5^. Ultra-Slow: *p*_*FDR*_ =2.32*×*10^−5^). In sum, 5-MeO-DMT makes low-frequency flow fields less recurrent, and less structured over time.

To test for changes in specific patterns, we classify singularities as either unstable node (sources), stable node (sinks), unstable focus (spiral-out), stable focus (spiral-in) or saddle waves (see Methods) (Fig.3g) and calculate their dynamical properties. We find that 5-MeO-DMT significantly reduces the average lifetime of every pattern of Delta and Slow waves, while engendering a mixed effect on Ultra-Slow waves, reducing the endurance of nodes but increasing the endurance of spirals and saddles (Fig.3h Delta max: S-Focus: *T* =-5.880, *p*_*FDR*_ =3.31*×*10^−4^, *BF* =371.185; Slow max: S-Node: *T* =-8.757, *p*_*FDR*_ =6.63*×*10^−5^, *BF* =1.20*×*10^4^; Ultra-Slow max: Saddle: *T* =5.463, *p*_*FDR*_ =5.42*×*10^−4^, *BF* =209.142) (See Supplementary Table.7 for extended statistics). Overall, these results show that 5-MeO-DMT disrupts canonical patterns of low-frequency flow.

**Figure 3.**
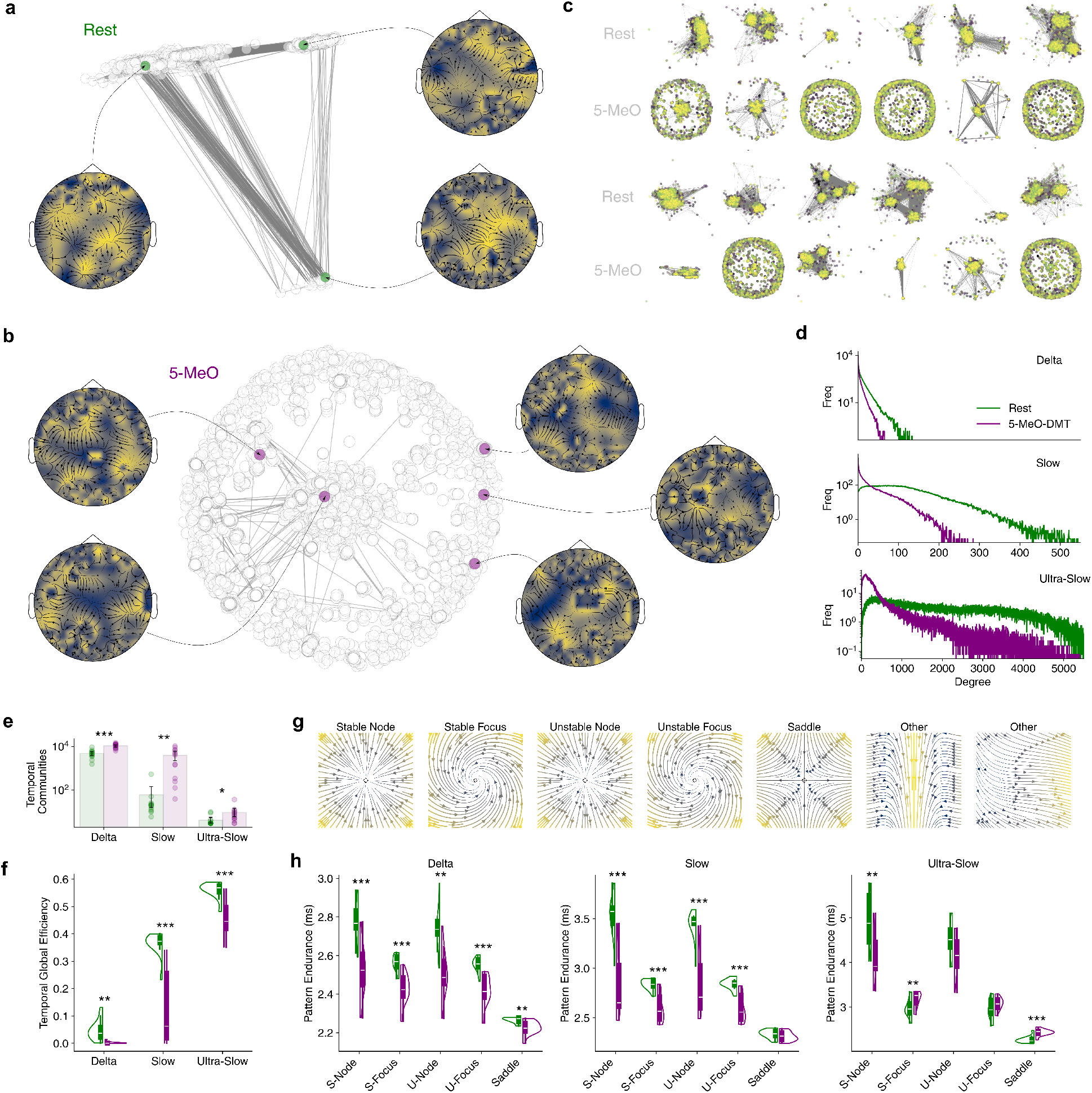
5-MeO-DMT dampens low-frequency field recurrence and pattern endurance. **a** Field recurrence network with example velocity fields from Mode 2 in the Rest condition over 15s of a representative participant. Plotted with communities determined by greedy modularity maximisation (25 modules). **b** Field recurrence network plot with same parameters for 5-MeO-DMT (1513 modules). **c** Field recurrence network with same parameters for remaining participants. **d** Mean histograms for degree distribution of field recurrence networks for each mode **e** Number of communities in recurrence networks for each mode. **f** Global efficiency for recurrence networks for each mode. **g**.Illustrations of canonical singularity patterns. **h** Average persistence of a flow pattern as the dominant pattern in the brain. S indicates a stable singularity, U indicates unstable. Sources (U-Nodes), sinks (S-Nodes), spiral-out (U-Focus), spiral-in (S-Focus) and saddles. (^*^,^**^,^***^ indicates p<.05,.01,.001 FDR-corrected Benjamini–Hochberg procedure)

### Broadband stability

We predicted that a potential consequence of these low-frequency alterations, and a potential cause of the structurally constrained experience induced by the drug, would be that broadband neural activity would be forced onto an unusual manifold, where large global shifts are prohibited. We expected that broadband signals would on average exhibit slower, more stable, low-dimensional behaviour, with increased energetic costs for major deviations.

First, we assessed the average decay time of the automutual information (*AMI*) function, a measure of the ‘intrinsic timescale’ of neural activity, and find that this doubles under 5-MeO-DMT from 221.9 ms *±* 72.6 to 495.3ms *±* 94.7 (Fig.4a; *p*_*FDR*_ =0.018), an effect robust to parameter choices (Supplementary Fig.6). Thus, 5-MeO-DMT makes broadband activity exhibit more regular trends and long-term dependence. We then compute the maximum Lyapunov exponent, a measure of the average separation rate of neighbouring points in phase space, and find a weak but significant reduction (*λ*_*max*_) (Fig.4b; *p*_*FDR*_ =0.046). Thus, 5-MeO-DMT makes cortical dynamics more stable by decreasing its sensitivity to initial conditions. We next tested whether cortical dynamics would be easier to explain via principal components analysis with singular value decomposition. We indeed show that the variance explained (the eigenvalue) is significantly greater for 5-MeO-DMT for each component (eigenvector) (Fig.1c; *PC*_1_ *p*_*FDR*_ =0.027), indicating that the brain under 5-MeO-DMT indeed exhibits more low-dimensional behaviour.

**Figure 4.**
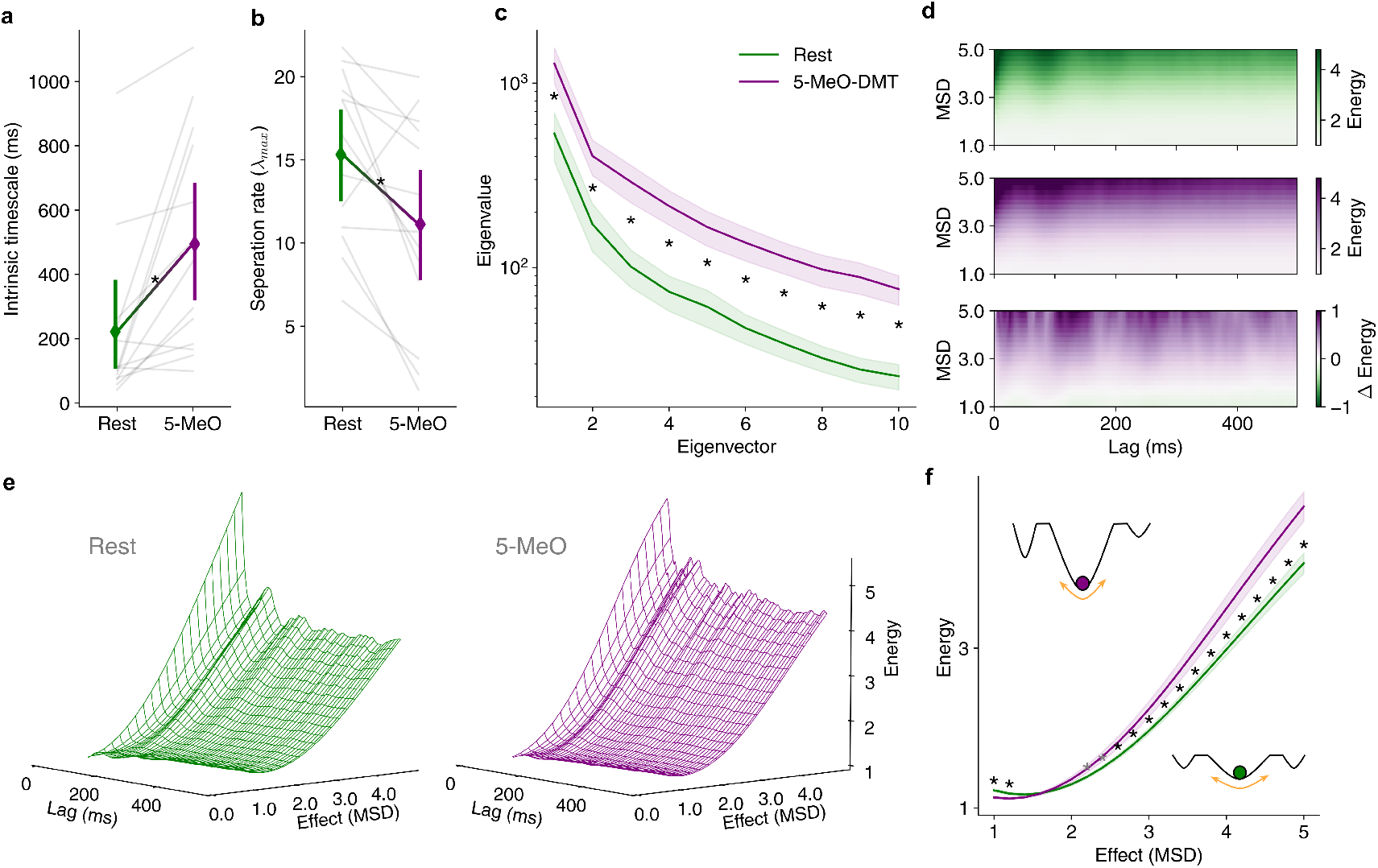
5-MeO-DMT pushes cortical dynamics toward a low-dimensional steady-state. **a** Average decay time of auto mutual information (*AMI*) function on broadband signals. **b** Average separation rate of neighbouring points in the phase space of broadband signals. **c** Eigenvalues for the first 10 eigenvectors from PCA of broadband signals. **d** 2D average energy landscape for Rest (top), 5-MeO-DMT (middle) and their difference (bottom). **e** 3D average energy landscape for Rest (left) and 5-MeO-DMT (right). **f** Energy required for each MSD bin averaged across all lags. ‘Energy well’ illustrations are included to depict energetic costs. (^*^ indicates p<.05 FDR-corrected)

Finally, we examine whether this slow, stable, low-dimensional broadband behaviour can be cashed out in the language of statistical mechanics, as alterations in the topography of a putative energy landscape of cortical dynamics. Here, energy represents the ability of the brain to move from one activity state to another at a particular timescale, and is operationalised as the inverse probability of a mean-squared displacement in global neural activity of a particular size as a function of time [49]. Given the hypothesis that 5-MeO-DMT pushes the brain into an unusual functional sub-space, we expected that 5-MeO-DMT would ‘protect’ the brain from major deviations in neural activity. Indeed, we find that 5-MeO-DMT significantly increases the energy requirement of large scale deviations (>3 MSD) (max *T* =5.527, *p*_*FDR*_ =0.001, *BF*_10_ =228.686, *d* =1.193), while also making minor shifts (<1.5 MSD) easier to attain (max *T* =-4.628, *p*_*FDR*_ =0.002, *BF*_10_ =62.859, *d* =1.307) Fig.4d. We also use non-linear least squares to fit *e*^*m·x*^ curves to our 2D landscape functions and find that 5-MeO-DMT significantly increases the slope of the estimated curve (*p* =1.58*×*10^−4^). Thus, 5-MeO-DMT constrains the global structure of neural dynamics.

## 3 Discussion

We demonstrate that 5-MeO-DMT, a short-acting psychedelic drug, produces a unique state in the human brain. This state is marked by amplified diffuse slow rhythmic activity, that coalesces into spatiotemporally disorganised and dismantled wave patterns unable to travel up and down the putative cortical hierarchy. Furthermore, we find that this disruption pushes broadband neural activity towards a low-dimensional steady-state, paralleling the subjective quality of the experience.

Our results undermine the hypothesis that the induction of diffuse high-amplitude slow oscillations are a universal signature of loss of consciousness. They do, however, provide evidence that augmented low-frequency oscillations are related to environmental disconnection, where subjective experience becomes independent of sensory variables imported from the external world [54, 73–75]. This result therefore reinforces the need for establishing more robust neuroscientific methods for discriminating conscious states. We offer the theoretical and empirical contribution that a fruitful strategy will be found by moving beyond one-size-fits-all univariate metrics such as oscillatory power, towards comprehensive frameworks that permit the detailed characterisation of complex spatiotemporal activity structures.

By tracking the extended spatiotemporal patterns of low-frequency flows of neural activity, we discovered crucial characteristics of the 5-MeO-DMT brain state that distinguish it from states of unconsciousness. Rather than triggering simple, coherent, fluid, persistent, recurrent propagating global waves like in anaesthesia [25, 27, 48], we found that 5-MeO-DMT induces complex, incoherent, viscous, fleeting, unique wave patterns. While these diverse local flows constrain the cortex-wide propagation, it may be the case that they enhance a form of regional collision-based distributed dynamical computation [40]. The 5-MeO-DMT state may therefore be marked by enhanced parallel, but not necessarily integrated, information processing [40]. Future work should investigate how spatiotemporal flow structures relate to taxonomies of multivariate information dynamics, including measures of integrated information [76].

We found that 5-MeO-DMT instigates more stable low-dimensional broadband behaviour with a decreased likelihood of major rapid global activity reconfiguration. These results are consistent with work in mice showing that cortical dynamics become less chaotic under 5-MeO-DMT [20]. This overall simplicity may reasonably occur as a consequence of, rather than in spite of, the complexity seen in low-frequency flows. The lack of collective spatiotemporal coordination of the low-frequency flow fields implies that cortical dynamics are sub-served by a more fragmented network architecture, which hinders the recruitment of multiple regions to effectively orchestrate simultaneous large global amplitude deviations. In short, a segregation across spatiotemporal scales occurs, deconstructing the brain’s canonical functional organisation.

Our results contest a number of key concepts in the developing literature on the neuroscience of serotonergic psychedelic drugs. First, whole-brain suppression of alpha power is considered a central feature of the psychedelic state [77]. However, we found that alpha oscillations are not robustly reduced across the cortex, with no deflation of the average power, and only right parietal-occipital electrodes reaching significance. Future studies should investigate how alterations in properties such as waveform shape and rhythmicity may be key. Second, psychedelics are thought to ‘liberate’ the bottom-up flow of neural activity [78, 79]. By contrast, we found a striking reduction in the probability that low-frequency flows travel in the anterior direction, as well as the posterior direction. Finally, psychedelics are thought to reduce the curvature of the energy landscape constraining neural activity, allowing more facile activity shifts, and higher dimensional dynamics [78, 80–82]. Yet, we found that 5-MeO-DMT steepened the brain’s energy landscape, increasing the barriers for major rapid activity shifts, and diminishing the dimensionality of neural dynamics. Our results therefore suggest that 5-MeO-DMT has a unique effect on the human brain compared to other classic tryptamine psychedelics.

Neurobiologically, we speculate that our results may represent the consequences of a unique shift in the balance of distinct thalamocortical subsystems. The presence of diffuse high-amplitude slow rhythmic activity, reduced communication through coherence, and approach toward a low-dimensional steady state, suggest a reduction in diffuse coupling in the brain. Non-specifically projecting thalamic structures, such as the mediodorsal and centromedian nuclei, may be relieved of their canonical tonic firing patterns orchestrating supragrangular cortical dynamics and global synchronisation patterns [83, 84]. Instead, cyclical bursting and quiescence of these nuclei may occur, resulting in functional deafferentiation, and cortical activity to become dominated by the intrinsic slow non-stationary bistability prescribed the anatomy of local cortical circuits [85–87]. However, as internal awareness and arousal is maintained, the level of overall thalamocortical resonance expected by wakefulness is likely maintained. Indeed, the presence of fleeting viscous and heteregenous local flow hints there could be markedly greater driving gain by structures with targeted projections, like the pulvinar and ventral lateral nucleus, disrupting local excitation-inhibitory balance and overwhelming wave pattern dynamics. We note that this general hypothesis is distinct from existing corticothalamic models of psychedelic action which posit that there is an indiscriminate reduction in thalamic gating [1, 88].

Acknowledging some limitations, the study suffers from a small number of participants with a high exclusion rate. This was due to movement artefacts, a common issue for human neuroimaging studies with psychedelic drugs. The control condition is a resting-state baseline, rather than a blinded placebo group, meaning that expectancy factors are uncontrolled for. A standardised single high-dose (12mg) dose was administered to each participant, and blood samples were not taken throughout, meaning that inter individual differences in pharmacokinetics are likely present. Methodologically, our analyses assume the cortex to be a relatively sparse flat surface, and therefore does not account for the high-spatial resolution, nor the gyri and sulci, inherent to the human brain. Subsequent studies should therefore investigate 5-MeO-DMT with high-density electrophysiological methods with effective cortical surface modelling. Lastly, the present work does not integrate neural measures with first-person reports. Accordingly, future work should combine neuroimaging with rigorous time-resolved measures of subjective experience, such that inferences about both can mutual constrain each other.

In conclusion, we present the first detailed neuroscientific investigation in humans of perhaps the most radically altered state of consciousness known. We report novel changes in the spatiotemporal organisation of low-frequency waves across the cortex, revealing a disruption in the congruity of the dynamical processes upon which the rest of cortical activity is built. This work not only underscores the need for a revitalised understanding of the role of slow waves in orchestrating subjective experience, but emphasises the need for more comprehensive spatiotemporal methods in neuroscience to better understand the full diversity of human brain states.

## 4 Methods

### Ethical statement

This study was conducted in accordance with the Declaration of Helsinki and approved by the University College London (UCL) Research Ethics Committee (ID: 19437/004). The study was conducted in collaboration with Tandava Retreat Centre (TRC), who provided trained facilitators and facilities. All participants provided informed consent via a secure online platform after reviewing comprehensive study information. Participation was voluntary and uncompensated. Participants were informed of their right to withdraw at any stage of the study without penalty and were provided with information about sources of support in case of any distress related to the study.

### Participants

Participants were recruited globally through the study advertisement on the F.I.V.E. (5-MeO-DMT Information & Vital Education) platform (www.five-meo.education), social media, news outlets, and word-of-mouth. Potential participants underwent a four-stage screening process: (1) TRC online screening, (2) TRC facilitator interview, (3) UCL online screening, and (4) UCL researcher interview (RM, GB). Exclusion criteria included: age <18 years, no prior 5-MeO-DMT experience, significant physical illness (e.g epilepsy, heart disease), psychiatric diagnoses, use of psychiatric medications of any kind, family history of psychosis, history of adverse reactions psychedelic drugs, and known physical movement under the influence of 5-MeO-DMT. Eligible participants (N = 32) were initially assigned to one of six three-day retreats, with assignments based on availability and balanced distribution. However, three participants withdrew before the retreats began. The final cohort (N = 29; 16 male, 13 female; mean age = 48.52 years, SD = 10.44, range = 34–75) was predominantly White/Caucasian (86.21%). The majority of participants held a Bachelor’s degree (58.62%) and identified as non-religious (93.10%). Mean lifetime 5-MeO-DMT use was 39.03 occasions (SD = 72.91, range = 1–300). Detailed participant characteristics are provided in Supplementary Table.1.

### Experimental procedures

The three-day retreat protocol included a preparation day consisting of orientation and participant briefing (Day 1), a dosing day for 5-MeO-DMT administration and primary data collection (Day 2), and an integration day featuring facilitator-guided group discussions (Day 3). This setup was consistent across all six retreats. Participants were instructed to refrain from consuming drugs, with the exception of nicotine containing products, for two weeks prior to dosing. On dosing day, participants fasted and abstained from caffeine. Individual dosing sessions commenced at 08:00 AM in 90-minute intervals, supervised by two lead facilitators. Each session began with a 7-minute baseline eye-closed resting-state EEG recorded with participants seated in a self-selected comfortable position conducive to minimal movement. Participants were instructed to remain relaxed but wakeful, and to alert experimenters if they felt like they may fall asleep. Participants then moved to a centrally located padded recliner, where they assumed a semi-supine position equipped with a standardised neck pillow and opaque eye mask. Synthetic 5-MeO-DMT (12 mg) was vaporised using an argon gas piston vaporiser (203-210°C, ∼120s heating). Following a standardised 2-3 minute relaxation exercise, participants inhaled the vapour in a single breath and were instructed to maximise breath retention before exhalation, as per protocols established during preparation. EEG recording began at the moment of completed inhalation, and continued for 20 minutes post-inhalation. The EEG system was a saline-based 64-channel ANT Neuro Waveguard Net connected to an NA-261 EEG amplifier, recorded at 500 Hz, with Ag electrode placement following the 10-20 system (reference: CPz, ground: AFz, impedance <20 kΩ). Ambient non-percussive music was played throughout the baseline and drug period, and a Koshi bell was rung when EEG recording was complete. Post-session, participants rested for a self-determined period.

### EEG preprocessing

Preprocessing was performed using custom scripts using MNE-Python [89].Participants were excluded from analysis if their physiological artifacts (ocular, muscular, respiratory, or cardiovascular) were substantial enough to necessitate the removal of over 50% of the data collected within the first 10 minutes post-drug administration (10 participants). Data was filtered using a finite impulse response (FIR) one-pass, zero-phase, non-causal bandpass filter at 0.1-50 Hz. Data was segmented into non-overlapping 3s epochs, producing 140 epochs for the baseline condition and 400 epochs for the drug condition. Electrolyte bridging was computed using the intrinsic Hjorth algorithm with a 16*µV* ^2^ cut-off [90, 91]. Participants with ≥1% of potential total bridges, or significant bridging that localises such that interpolation would depend on sensors which are also bridged, were excluded from further analysis (1 participant). For those with <1%, spherical spline interpolation was performed via the generation of virtual channels between bridged electrodes (Rest: 3.684 *±* 2.076, 5-MeO-DMT: 1.158 *±* 0.785 bridges). Sensor time series were manually inspected to identify bad channels, participants with >20 bad channels were excluded from further analysis (0 participants), and those with <20 bad channels had these interpolated from surrounding sensors (Rest: 7.579 *±* 1.049, 5-MeO-DMT: 7.368 *±* 1.06 channel). Extended infomax independent components analysis (ICA) was performed with 40 components, and those manually deemed to be equipment noise or physiological artefacts were zeroed out and excluded from sensor signal reconstruction (Rest: 4.842 *±* 0.766, 5-MeO-DMT: 5.053 *±* 0.89 components). Epochs corrupted by noise or physiological arefacts were manually removed (Rest: 3.611 *±* 2.267, 5-MeO-DMT: 38.105 *±* 10.718 epochs). Data was baseline corrected and rereferenced to an average reference. This resulted in 19 subjects, 13 of which had an artifact-free continuous minutes at the peak of the drug’s effects.

### Empirical Mode Decomposition

A central challenge in time series analysis is creating robust representations of the frequency content of complex signals. The standard approach in neuroscience for doing so is based on the Fourier Transform (FT). The FT is a powerful and versatile technique that seeks to represent a signal as a combination of strictly linear sinusoidal basis functions. However, brain rhythms are noisy, exhibit distinctly non-sinusoidal waveforms, and enter into transient bursting behaviour [65–70]. This presents a challenge for the FT, since it cannot fully and directly represent the non-sinusoidal forms at their intrinsic frequency, instead assuming them to be partly the result of a higher frequency harmonic [92]. This is problematic for neuroscience, since the canonical electrophysiological frequency bands are all harmonics of a lower band, and the power in each specific band is a variable of interest in its own right for distinguishing different brain states. This problem is exacerbated by the fact that brain signals comprise a myriad of multiplexing oscillations, each with their own harmonics. Furthermore, the linear basis functions of the FT are static, meaning that in order to characterise evolving brain activity analysts must use a windowing technique that inherently blurs the ongoing dynamics.

An elegant solution to these problems comes from Empirical Mode Decomposition (EMD), which permits the automatic decomposition of signals into a finite number of distinct dynamic non-linear oscillatory modes, known as Intrinsic Mode Functions (IMFs) [93]. EMD sifts the IMFs out of a signal, from fastest to slowest, in an iterative process of peak-trough detection, amplitude envelope interpolation and subtraction. Here, we use the recently developed iterated-masking EMD (itEMD), as it has been shown to be an automated solution to the mode mixing problem [72, 94, 95].

We then compute the instantaneous phase, frequency and amplitude of each IMF via the Normalized Hilbert-Huang transform (HHT) [96, 97], with the signal’s time-resolved power spectrum being an instantaneous amplitude weighted representation of frequency content [98]. Finally, to investigate the specific frequency content of amplitude modulations in our signals, we perform a second level sift on the instantaneous amplitudes of each IMF, producing a set of higher-order IMFs which represent the frequency content (amplitude modulation frequency) of dynamical changes in oscillatory power (carrier frequency) [97]. Carrier frequencies are placed into 50 linearly spaced histogram bins 0.5-4Hz and amplitude-modulation frequencies are placed into 50 linearly spaced bins 8-50Hz, producing the holospectrum.

### Velocity fields

Velocity vector fields are mathematical tools from physical theories of fluid dynamics that include descriptions of vortices and eddies in collective fluid motion [99]. Recently, velocity fields have been appropriated in neuroscience to characterise the structure of large-scale propagating patterns of neural activity, adapting techniques from optical flow estimation in computer science [36, 100].

First, the instantaneous amplitude of an IMF is pushed onto a 32×32 scalp grid, using radial basis function interpolation and a multidimensional gaussian kernel of width 1 S.D, creating the sequence *ϕ*(*x, y, t*). The velocity field **v**_*ϕ*_(*x, y, t*) = (**u**(*x, y, t*), **v**(*x, y, t*)), of element-wise x and y-magnitudes, is then simply recovered from the spatial and temporal derivatives of the amplitude via solving the constancy equation [27]:

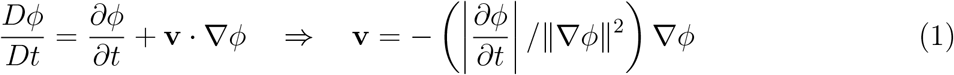

### Order parameters

#### Synchrony

As is done with the classical Kuramoto order parameter [101], we compute the synchrony of instantaneous phases *θ* as:

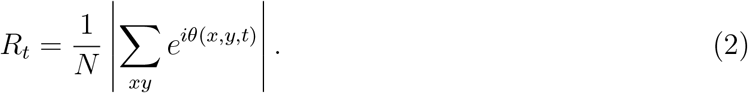

Coherence is then the mean of this parameter across time, and metastability is the variance across time [102]. We compute this both globally, and locally across radii ranging from 1.750 cm to 7.875 cm in increments of 0.875. The minimum corresponds to one grid element and anything significantly larger than the maximum would entail areas that are predominantly outside of the scalp.

#### Normalised Velocity

To characterise the collective motion of the field, we calculate the norm of the sum of the velocities divided by the sum of the norm of the velocities:

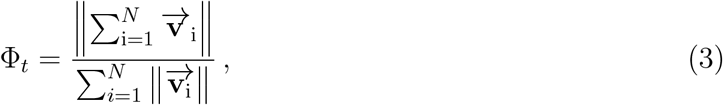

where *N* is the number of spatial elements in the field and 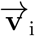 is the velocity the i-th element. If Φ_*t*_ is close to 1, the vectors in the field are aligned and there is collective motion in a particular direction [41, 103].

#### Heterogeneity

To characterise the diversity of flows in the field we compute the heterogeneity [48], as the mean over time of the standard deviation of wave propagation speeds. More formally:

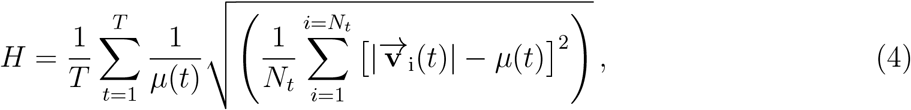

where *T* is the total time, *N*_*t*_ the number of vectors at time t, and 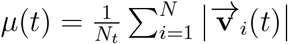 the average speed at time t.

### Complex wave patterns

Complex spatiotemporal waves are organised around critical points of stationarity [104]. These singularities are of two main flavours, sources where flow arises, and sinks where flow gathers. First, we compute the divergence of the velocity field as the sum of the partial derivatives of each field component 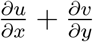. Singularities were then identified as extremal nested closed contours divergence [36]. Contours were run over 10 levels equally spaced between the 1st and 99th percentile of the divergence values [27]. From this we compute the centroid of the singularity, the area of the contour occupied by the singularity, and the strength of the singularity as the magnitude of velocity divergence.

This allows us to calculate viscosity of flow, which is taken simply as the product of the area and divergence of the singularities [27], giving us a cm^2^s^−1^ unit, more formally:

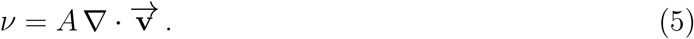

Lower values indicate that waves can flow across space with greater ease, whereas higher values indicate their distributed diffusion is restricted.

The nature of the singularities is then further characterised by the values of the trace (*τ*) and the determinant (Δ) of the Jacobian *J* [36], bilinearly interpolated from the neighbouring grid elements on the scalp, which correspond to the divergence and curl of the field respectively:

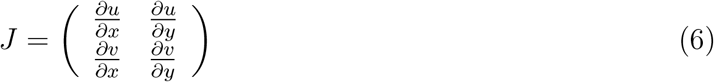

Unstable singularities (sources) have *τ <* 0 and stable singularities (sinks) have *τ >* 0. Nodes have Δ *>* 0 and *τ* ^2^ *>* 4Δ, focuses have Δ *>* 0 and *τ* ^2^ *<* 4Δ. Stable focuses are spiral-in waves, and unstable focuses are spiral-out waves. Lastly, saddles are characterised by Δ *<* 0, and are usually formed by interactions between other waves [36].

### Velocity field recurrence

In order to characterise the recurrence of velocity fields we calculated their alignment matrix [30] via their mutual information (*MI*):

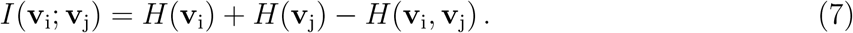

Where *H* is the classic Shannon entropy function. We then shuffle the velocities spatially, compute the alignment matrix over 100 permutations and take the maximum *MI* value from each permutation and chose the 99th percentile of these values as a threshold for our empirical matrix. We treat this as the binary adjacency matrix *A* of a temporal graph, and characterise its structure [105].

### Recurrence network topology

We first cluster nodes using Clauset-Newman-Moore greedy modularity maximization [106], which finds the network partition that maximises the generalised modularity *Q*:

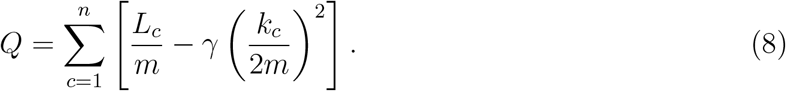

Where the sum iterates over all communities *c, L*_*c*_ is the number of intra-community edges, *m* is the total number of edges, *k*_*c*_ is the sum of the degrees in the community, and *γ* is the resolution parameter which we set to 1 [107].

We also characterise the compactness of this recurrence network by computing the average of the inverse distances between nodes, i.e. its global efficiency:

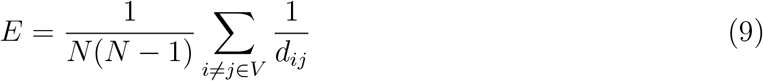

, where *d*_*ij*_ is the distance between the i-th and j-th nodes. In this case, higher global efficiency indicates that the network is compact and velocity fields are robust over time, whereas a lower global efficiency indicates that velocity fields exhibit less recurrence.

### Intrinsic timescales

To identify the intrinsic timescale of the cortex we compute the average decay time of the auto-mutual information (*AMI*) function. That is, we find the mean lag at which the mutual information between neural signals and a delayed copy of themselves reaches a stable minima. To do so, we fit a straight line to the last third of the *AMI* function, and find the first time point that falls below the *AMI* value equal to the line’s y-intercept. Longer intrinsic timescales indicate that a time series contains greater information about its own future, and shorter intrinsic timescales indicate that a timeseries is less history dependent [108].

### Separation rate

To characterise the dynamical sensitivity of the system, we characterise the rate of divergence of similar points in phase space with the maximum Lyapunov exponent:

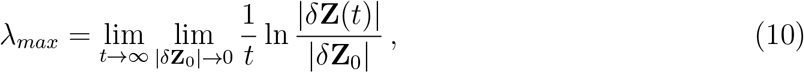

which we derive via Rosenstein’s algorithm [109]. Here *δ***Z**(*t*) is the trajectory separation at time *t*. Briefly, the broadband amplitude envelope is produced by the Hilbert transform and its dynamics are reconstructed using Taken’s embedding. The nearest neighbour of each time point is identified using the Euclidian distance, and their trajectories tracked. The separation rate is then the exponent of the curve of the mean logarithmic distance between these trajectories as a function of time, normalised by the sampling rate.

### Energy landscape

To characterise the shifts in the ability of the brain to orchestrate rapid global broadband reconfigurations, we follow previous work [49, 110, 111], and characterise the energy landscape of neural dynamics through the statistical inference of the probability of an effect of a particular size. Effect is operationalised as mean-squared-displacement (MSD) in the Fisher z-scored amplitude envelope of broadband signals across the cortex:

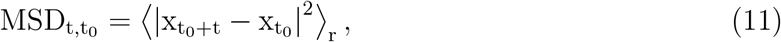

where *r* is the number of electrodes. We compute the MSD with maximum lags of 1s, with 200 equally spaced initialisation points. As is done in statistical mechanics, we can then compute the energy *E* of the electrophysiological attractor basin via the natural logarithm of the inverse probability, estimated via Gaussian kernel density estimation K. Following [111], we compute this over MSD values between 0 and 5.

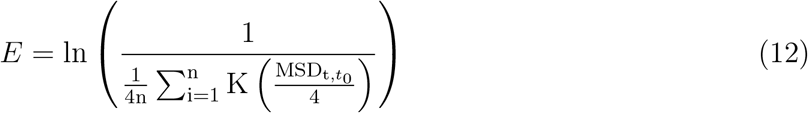

Thus, a highly probable relative change in neural activity corresponds to a putatively low energy requirement, and vice versa.

## 5 Data availability

Data will be made publicly available upon acceptance of the manuscript for publication.

## 6 Code availability

Code will be made publicly available upon acceptance of the manuscript for publication.

## 7 Acknowledgements

We are grateful to Tandava Retreats for their generous support in assisting the data collection process, specifically Victoria Wueschner and Joel Brierre for their pivotal roles in facilitating the 5-MeO-DMT sessions. Additional thanks go to Luis Fabian Rodriguez, Otto Maier, James Sanders, Milly Sellers and George Deane for providing support during data collection. We extend our deepest thanks to the courageous participants who volunteered for this study. Lastly, we thank supporters of the crowdfunding campaign for helping secure the funding required to perform this study. G.B is supported by the Leverhulme Trust. R.G.M is supported by the Wellcome Trust. J.I.S. is supported by Wellcome Leap.

## 8 Contributions

G.B; Conceptualization, Investigation, Methodology, Software, Data curation, Formal analysis, Supervision, Project administration, Writing. R.G.M; Conceptualization, Investigation, Project Administration, Writing – Review and Editing. M.S.F; Methodology, Software, Writing. A.L; Writing – Review and Editing. S.K.K; Resources, Supervision, Writing – Review and Editing. P.A.M.M; Supervision, Writing – Review and Editing. J.I.S; Supervision, Writing – Review and Editing.

