## Supplementary Material for "Complex slow waves radically reorganise human brain dynamics under 5-MeO-DMT"

October 4, 2024



This work is shared under a CC BY license unless otherwise noted

Table 1: **Participant information**

| Sub | Epoch | Cont | Age | Gender | Ethnicity | Education | Religion | Lifetime Use |
|---|---|---|---|---|---|---|---|---|
| 1 | 0 | 0 | 34 | Female | Asian | Bachelor's | No | 5 |
| 2 | 1 | 1 | 68 | Female | Latino | Bachelor's | No | 25 |
| 3 | 1 | 1 | 47 | Male | Latino | Bachelor's | No | 2 |
| 4 | 1 | 0 | 55 | Female | White | Bachelor's | No | 20 |
| 5 | 1 | 1 | 46 | Female | White | Bachelor's | No | 50 |
| 6 | 1 | 0 | 47 | Female | White | Bachelor's | No | 15 |
| 7 | 1 | 0 | 54 | Female | White | Bachelor's | No | 6 |
| 8 | 0 | 0 | 41 | Female | White | Bachelor's | Yes | 4 |
| 9 | 0 | 0 | 41 | Female | White | Bachelor's | No | 3 |
| 10 | 0 | 0 | 54 | Female | White | Bachelor's | No | 1 |
| 11 | 1 | 1 | 50 | Male | White | Bachelor's | No | 12 |
| 12 | 1 | 1 | 65 | Male | White | Bachelor's | No | 10 |
| 13 | 1 | 1 | 42 | Male | White | Bachelor's | No | 100 |
| 14 | 1 | 0 | 40 | Male | White | Bachelor's | Unsure | 5 |
| 15 | 1 | 1 | 35 | Male | White | Bachelor's | No | 40 |
| 16 | 1 | 1 | 60 | Male | White | Bachelor's | No | 3 |
| 17 | 1 | 0 | 34 | Male | White | Bachelor's | No | 2 |
| 18 | 0 | 0 | 49 | Male | Latino | $H$igh School | No | 20 |
| 19 | 0 | 0 | 37 | Female | White | Master's | No | 5 |
| 20 | 0 | 0 | 62 | Female | White | Master's | No | 5 |
| 21 | 1 | 1 | 52 | Female | White | Master's | No | 2 |
| 22 | 0 | 0 | 43 | Female | White | Master's | No | 100 |
| 23 | 0 | 0 | 41 | Male | White | Master's | No | 5 |
| 24 | 1 | 1 | 45 | Male | White | Master's | No | 1 |
| 25 | 1 | 1 | 36 | Male | White | Ph.D. or above | No | 2 |
| 26 | 1 | 1 | 51 | Male | White | Ph.D. or above | No | 6 |
| 27 | 0 | 0 | 75 | Male | White | Not disclosed | No | 233 |
| 28 | 1 | 0 | 49 | Male | White | Trade School | No | 150 |
| 29 | 1 | 1 | 54 | Male | White | Trade School | No | 300 |



This work is shared under a CC BY license unless otherwise noted

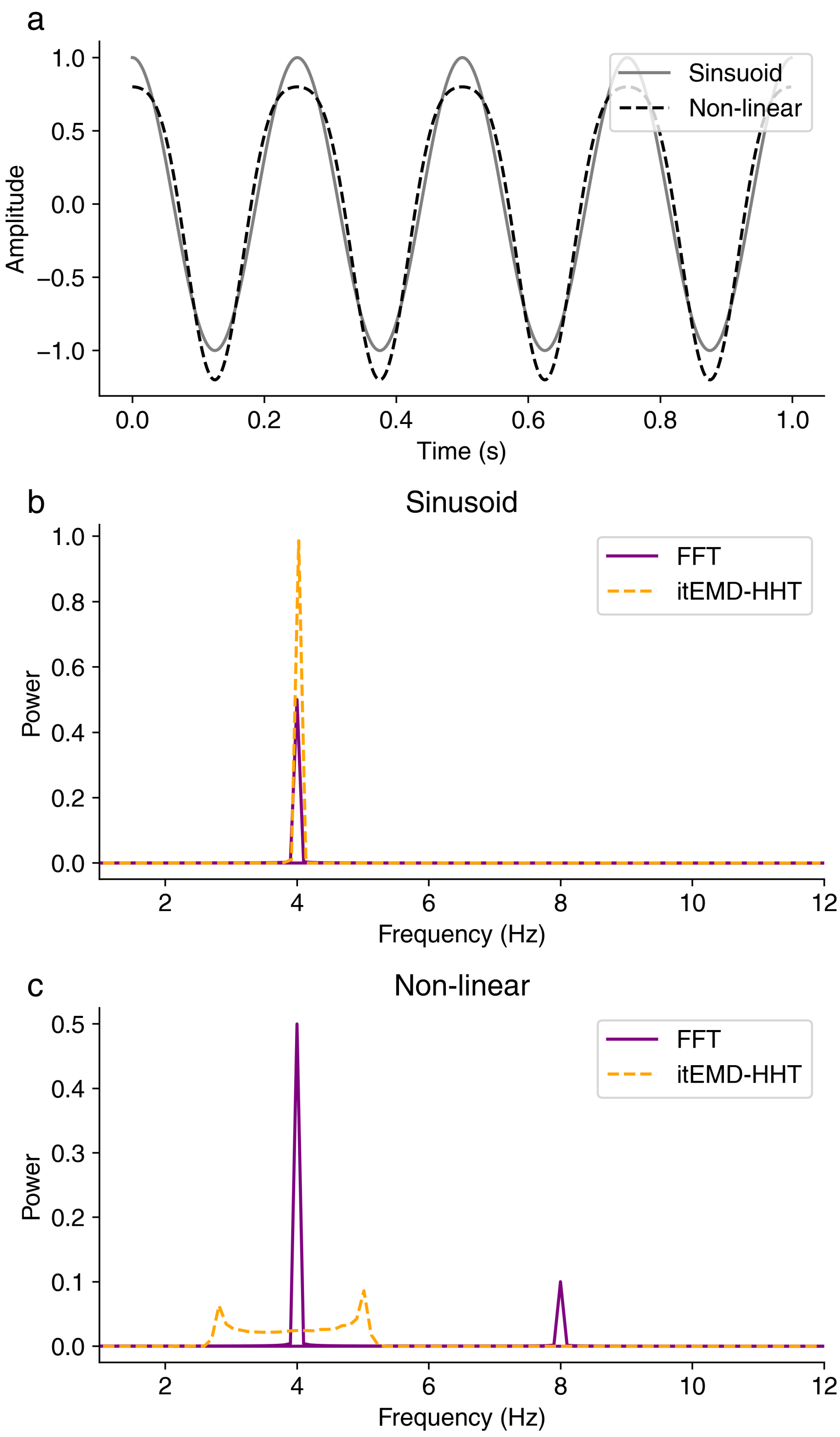


Figure 1: **FFT confabulates harmonic power for non-linear oscillations** **a** Linear oscillation $f(t) = \cos(2\pi ft)$, and non-linear $f(t) = \cos(2\pi ft) + \frac{1}{4}\cos(4\pi ft - \pi)$ oscillation with frequency of 4Hz. **b** Fast Fourier transform (FFT) power spectrum for the two signals **c** Iterated-masking empirical mode decomposition (it-EMD) Hilbert-Huang transform (HHT) power spectrum for the two signals



This work is shared under a CC BY license unless otherwise noted

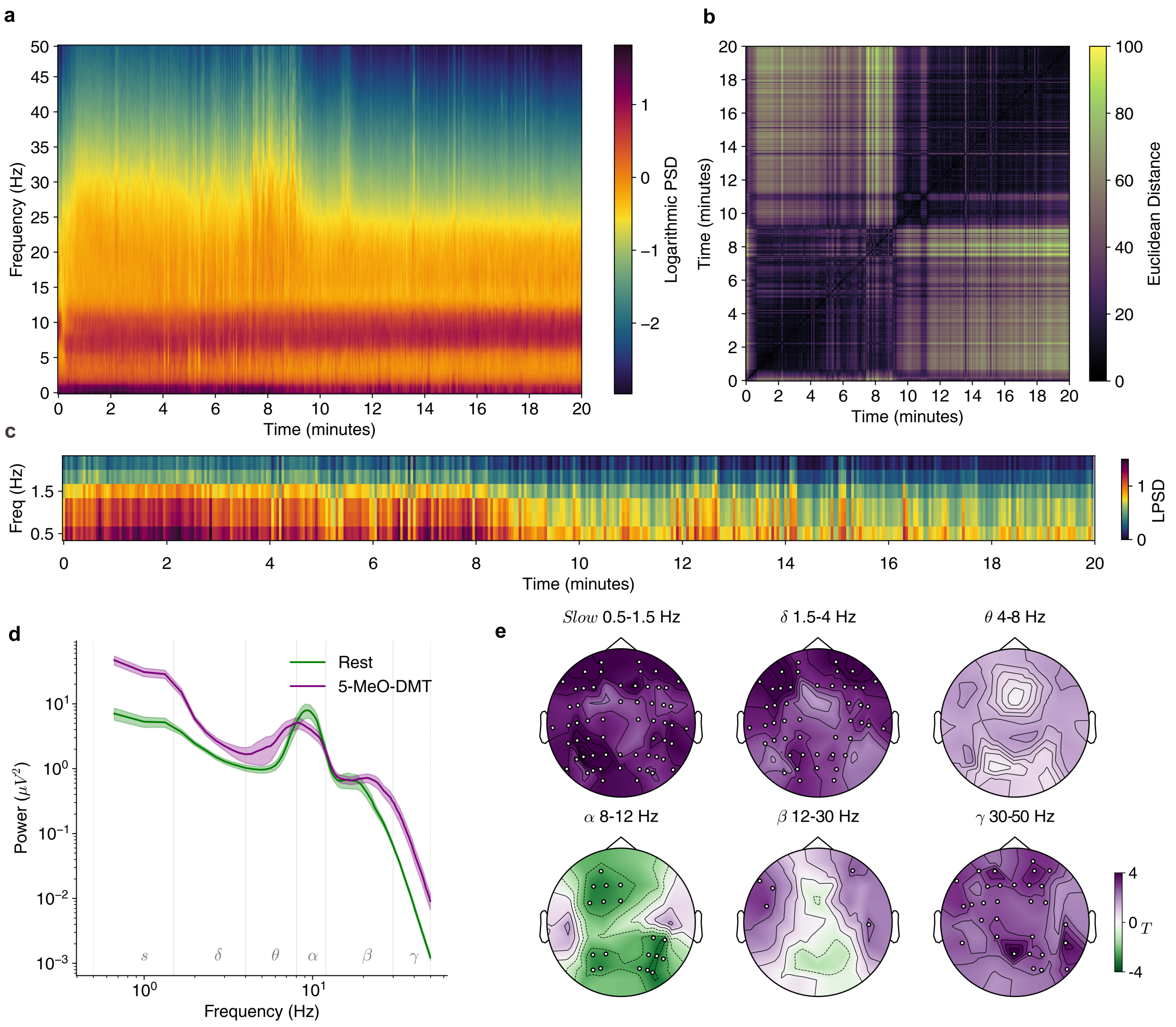


Figure 2: **Repetition of main manuscript Fig.1 EMD results with FFT a** Average time-frequency power spectral density (PSD) for the 20 minutes post-inhalation of 5-MeO-DMT, derived via multi-taper FFT with discrete prolate spheroidal sequence tapers. **b** Euclidean distance matrix for average spectra vectors over time. **c** Time-averaged power spectra for eyes-closed baseline period and estimated peak 5-MeO-DMT at 1.5-2.5 minutes (mean ± SEM). **d** Focused low-frequency spectrogram. **e** Scalp topographic statistical maps for T-test comparing the two conditions per frequency band and electrode, with electrodes representing significant differences ($p<.05$ FDR-corrected Benjamini–Hochberg procedure). Purple indicates increases under 5-MeO-DMT, green indicates reductions.

Table 2: **Kuramoto order parameter statistics**

| Measure | Mode | $T$ | $p$ | $d$ | $BF_{10}$ | $p_{FDR}$ |
|---|---|---|---|---|---|---|
| $C$ | $\delta$ | -3.174 | 0.008 | 1.432 | 6.948 | 0.010 |
| $C$ | S | -7.496 | 7.28e-06 | 2.784 | 2.89e+03 | 2.07e-05 |

Continued on next page



This work is shared under a CC BY license unless otherwise noted

Table 2: **Kuramoto order parameter statistics**

| Measure | Mode | $T$ | $p$ | $d$ | $BF_{10}$ | $p_{FDR}$ |
|---|---|---|---|---|---|---|
| $C$ | U-S | -11.223 | 1.02e-07 | 3.151 | 1.33e+05 | 8.73e-07 |
| $K$ | $\delta$ | -1.923 | 0.079 | 0.872 | 1.147 | 0.080 |
| $K$ | S | -4.885 | 3.75e-04 | 1.766 | 91.633 | 5.96e-04 |
| $K$ | U-S | -11.856 | 5.53e-08 | 3.395 | 2.30e+05 | 8.73e-07 |
| $\overline{C}_l$ 2 | $\delta$ | -3.103 | 0.009 | 1.404 | 6.238 | 0.011 |
| $\overline{C}_l$ 2 | S | -6.349 | 3.67e-05 | 2.602 | 6.93e+02 | 7.62e-05 |
| $\overline{C}_l$ 2 | U-S | -10.067 | 3.33e-07 | 3.122 | 4.55e+04 | 1.80e-06 |
| $\overline{K}_l$ 2 | $\delta$ | -1.606 | 0.134 | 0.705 | 0.777 | 0.134 |
| $\overline{K}_l$ 2 | S | -4.717 | 5.00e-04 | 2.000 | 71.673 | 7.71e-04 |
| $\overline{K}_l$ 2 | U-S | -3.856 | 0.002 | 1.493 | 19.690 | 0.003 |
| $\overline{C}_l$ 3 | $\delta$ | -3.265 | 0.007 | 1.480 | 7.981 | 0.008 |
| $\overline{C}_l$ 3 | S | -6.708 | 2.17e-05 | 2.697 | 1.10e+03 | 4.88e-05 |
| $\overline{C}_l$ 3 | U-S | -10.685 | 1.74e-07 | 3.184 | 8.16e+04 | 1.04e-06 |
| $\overline{K}_l$ 3 | $\delta$ | -2.386 | 0.034 | 1.082 | 2.158 | 0.036 |
| $\overline{K}_l$ 3 | S | -5.743 | 9.26e-05 | 2.391 | 3.08e+02 | 1.52e-04 |
| $\overline{K}_l$ 3 | U-S | -6.577 | 2.62e-05 | 2.426 | 9.31e+02 | 5.67e-05 |
| $\overline{C}_l$ 4 | $\delta$ | -3.364 | 0.006 | 1.521 | 9.288 | 0.007 |
| $\overline{C}_l$ 4 | S | -6.986 | 1.46e-05 | 2.761 | 1.56e+03 | 3.43e-05 |
| $\overline{C}_l$ 4 | U-S | -10.980 | 1.29e-07 | 3.203 | 1.07e+05 | 8.73e-07 |
| $\overline{K}_l$ 4 | $\delta$ | -2.577 | 0.024 | 1.176 | 2.839 | 0.027 |
| $\overline{K}_l$ 4 | S | -6.068 | 5.60e-05 | 2.458 | 4.78e+02 | 1.03e-04 |
| $\overline{K}_l$ 4 | U-S | -7.397 | 8.31e-06 | 2.625 | 2.57e+03 | 2.15e-05 |
| $\overline{C}_l$ 5 | $\delta$ | -3.430 | 0.005 | 1.546 | 10.261 | 0.007 |
| $\overline{C}_l$ 5 | S | -7.213 | 1.07e-05 | 2.802 | 2.06e+03 | 2.62e-05 |
| $\overline{C}_l$ 5 | U-S | -11.095 | 1.15e-07 | 3.203 | 1.18e+05 | 8.73e-07 |
| $\overline{K}_l$ 5 | $\delta$ | -2.611 | 0.023 | 1.196 | 2.986 | 0.027 |
| $\overline{K}_l$ 5 | S | -6.207 | 4.54e-05 | 2.459 | 5.75e+02 | 9.08e-05 |
| $\overline{K}_l$ 5 | U-S | -7.998 | 3.77e-06 | 2.745 | 5.19e+03 | 1.36e-05 |
| $\overline{C}_l$ 6 | $\delta$ | -3.464 | 0.005 | 1.557 | 10.820 | 0.006 |
| $\overline{C}_l$ 6 | S | -7.392 | 8.37e-06 | 2.834 | 2.56e+03 | 2.15e-05 |
| $\overline{C}_l$ 6 | U-S | -11.087 | 1.16e-07 | 3.193 | 1.18e+05 | 8.73e-07 |
| $\overline{K}_l$ 6 | $\delta$ | -2.588 | 0.024 | 1.187 | 2.889 | 0.027 |
| $\overline{K}_l$ 6 | S | -6.156 | 4.90e-05 | 2.399 | 5.38e+02 | 9.45e-05 |
| $\overline{K}_l$ 6 | U-S | -8.353 | 2.41e-06 | 2.808 | 7.74e+03 | 9.29e-06 |
| $\overline{C}_l$ 7 | $\delta$ | -3.478 | 0.005 | 1.559 | 11.051 | 0.006 |
| $\overline{C}_l$ 7 | S | -7.512 | 7.12e-06 | 2.853 | 2.95e+03 | 2.07e-05 |
| $\overline{C}_l$ 7 | U-S | -11.078 | 1.17e-07 | 3.185 | 1.17e+05 | 8.73e-07 |
| $\overline{K}_l$ 7 | $\delta$ | -2.544 | 0.026 | 1.166 | 2.710 | 0.028 |
| $\overline{K}_l$ 7 | S | -6.055 | 5.72e-05 | 2.318 | 4.70e+02 | 1.03e-04 |
| $\overline{K}_l$ 7 | U-S | -8.450 | 2.14e-06 | 2.815 | 8.61e+03 | 8.96e-06 |
| $\overline{C}_l$ 8 | $\delta$ | -3.483 | 0.005 | 1.558 | 11.137 | 0.006 |

Continued on next page



This work is shared under a CC BY license unless otherwise noted

Table 2: **Kuramoto order parameter statistics**

| Measure | Mode | $T$ | $p$ | $d$ | $BF_{10}$ | $p_{FDR}$ |
|---|---|---|---|---|---|---|
| $\overline{C}_l$ 8 | S | -7.602 | 6.32e-06 | 2.867 | 3.28e+03 | 2.01e-05 |
| $\overline{C}_l$ 8 | U-S | -11.077 | 1.17e-07 | 3.180 | 1.16e+05 | 8.73e-07 |
| $\overline{K}_l$ 8 | $\delta$ | -2.504 | 0.028 | 1.146 | 2.555 | 0.030 |
| $\overline{K}_l$ 8 | S | -5.984 | 6.37e-05 | 2.249 | 4.28e+02 | 1.11e-04 |
| $\overline{K}_l$ 8 | U-S | -8.443 | 2.16e-06 | 2.785 | 8.54e+03 | 8.96e-06 |
| $\overline{C}_l$ 9 | $\delta$ | -3.474 | 0.005 | 1.552 | 10.978 | 0.006 |
| $\overline{C}_l$ 9 | S | -7.651 | 5.92e-06 | 2.874 | 3.48e+03 | 2.00e-05 |
| $\overline{C}_l$ 9 | U-S | -11.075 | 1.18e-07 | 3.176 | 1.16e+05 | 8.73e-07 |
| $\overline{K}_l$ 9 | $\delta$ | -2.462 | 0.030 | 1.125 | 2.404 | 0.032 |
| $\overline{K}_l$ 9 | S | -5.923 | 6.99e-05 | 2.196 | 3.94e+02 | 1.18e-04 |
| $\overline{K}_l$ 9 | U-S | -8.502 | 2.01e-06 | 2.765 | 9.11e+03 | 8.96e-06 |

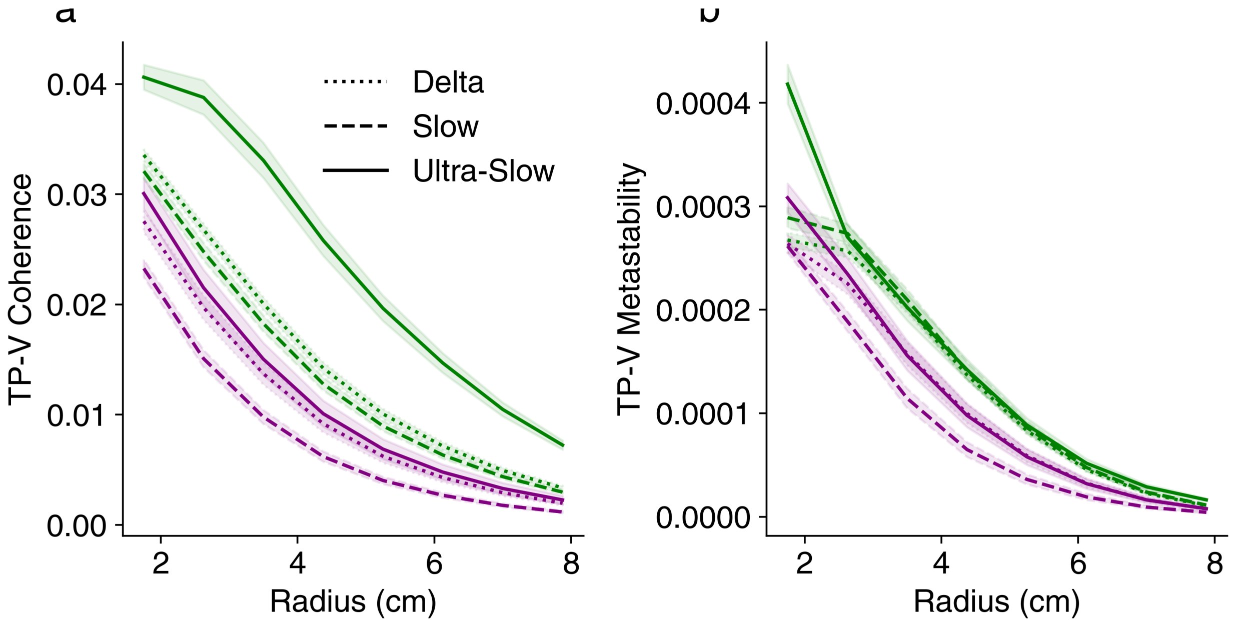


Figure 3: **5-MeO-DMT reduces the hierarchy of local synchronisation properties a** Reduction in variance across the scalp of local coherence over a range of radii (mean ± SEM) **b** Reduction in variance across the scalp of local metastability over a range of radii(mean ± SEM)

Table 3: **Velocity field order parameter statistics**

| Measure | Mode | $T$ | $p$ | $d$ | $BF_{10}$ | $p_{FDR}$ |
|---|---|---|---|---|---|---|
| $\overline{\Phi}$ | $\delta$ | -3.846 | 0.002 | 1.557 | 19.386 | 0.003 |
| $\overline{\Phi}$ | S | -7.426 | 7.99e-06 | 2.186 | 2.66e+03 | 7.19e-05 |
| $\overline{\Phi}$ | U-S | 2.541 | 0.026 | 0.830 | 2.697 | 0.026 |
| $H$ | $\delta$ | 4.513 | 7.10e-04 | 1.810 | 53.042 | 0.001 |

Continued on next page



This work is shared under a CC BY license unless otherwise noted

Table 3: **Velocity field order parameter statistics**

| Measure | Mode | $T$ | $p$ | $d$ | $BF_{10}$ | $p_{FDR}$ |
|---|---|---|---|---|---|---|
| $H$ | S | 6.203 | 4.57e-05 | 2.189 | 5.72e+02 | 1.49e-04 |
| $H$ | U-S | 5.959 | 6.62e-05 | 1.829 | 4.13e+02 | 1.49e-04 |
| Speed | $\delta$ | 4.153 | 0.001 | 1.669 | 30.889 | 0.002 |
| Speed | S | 6.037 | 5.87e-05 | 2.121 | 4.59e+02 | 1.49e-04 |
| Speed | U-S | 5.461 | 1.45e-04 | 1.729 | 2.09e+02 | 2.61e-04 |

Table 4: **Wave direction statistics**

| Mode | Angle | $T$ | $p$ | $d$ | $BF_{10}$ | $p_{FDR}$ |
|---|---|---|---|---|---|---|
| $\delta$ | -3.142 | -2.692 | 0.020 | 0.811 | 3.365 | 0.066 |
| $\delta$ | -3.013 | 0.738 | 0.475 | 0.281 | 0.352 | 0.603 |
| $\delta$ | -2.885 | 1.617 | 0.132 | 0.572 | 0.786 | 0.230 |
| $\delta$ | -2.757 | 2.377 | 0.035 | 0.770 | 2.131 | 0.097 |
| $\delta$ | -2.629 | 2.349 | 0.037 | 0.870 | 2.046 | 0.100 |
| $\delta$ | -2.500 | 2.232 | 0.045 | 0.902 | 1.738 | 0.113 |
| $\delta$ | -2.372 | 1.194 | 0.256 | 0.470 | 0.503 | 0.379 |
| $\delta$ | -2.244 | 3.357 | 0.006 | 1.365 | 9.186 | 0.031 |
| $\delta$ | -2.116 | 3.382 | 0.005 | 1.374 | 9.540 | 0.030 |
| $\delta$ | -1.988 | 2.081 | 0.060 | 0.803 | 1.413 | 0.136 |
| $\delta$ | -1.859 | 1.444 | 0.174 | 0.451 | 0.648 | 0.287 |
| $\delta$ | -1.731 | -1.312 | 0.214 | 0.426 | 0.564 | 0.328 |
| $\delta$ | -1.603 | -2.289 | 0.041 | 0.914 | 1.881 | 0.108 |
| $\delta$ | -1.475 | -2.659 | 0.021 | 1.038 | 3.204 | 0.066 |
| $\delta$ | -1.346 | -3.379 | 0.005 | 1.155 | 9.500 | 0.030 |
| $\delta$ | -1.218 | -1.001 | 0.337 | 0.399 | 0.425 | 0.459 |
| $\delta$ | -1.090 | 0.036 | 0.972 | 0.015 | 0.278 | 0.972 |
| $\delta$ | -0.962 | 0.308 | 0.763 | 0.129 | 0.290 | 0.830 |
| $\delta$ | -0.833 | -1.476 | 0.166 | 0.611 | 0.670 | 0.279 |
| $\delta$ | -0.705 | -0.057 | 0.956 | 0.022 | 0.279 | 0.962 |
| $\delta$ | -0.577 | 1.409 | 0.184 | 0.521 | 0.624 | 0.297 |
| $\delta$ | -0.449 | 1.456 | 0.171 | 0.478 | 0.656 | 0.285 |
| $\delta$ | -0.321 | 1.054 | 0.312 | 0.350 | 0.444 | 0.438 |
| $\delta$ | -0.192 | 0.328 | 0.748 | 0.100 | 0.292 | 0.823 |
| $\delta$ | -0.064 | -1.919 | 0.079 | 0.654 | 1.141 | 0.169 |
| $\delta$ | 0.064 | -2.250 | 0.044 | 0.776 | 1.782 | 0.113 |
| $\delta$ | 0.192 | -0.426 | 0.677 | 0.144 | 0.301 | 0.776 |
| $\delta$ | 0.321 | 0.138 | 0.893 | 0.039 | 0.281 | 0.905 |
| $\delta$ | 0.449 | 0.219 | 0.831 | 0.060 | 0.284 | 0.877 |
| $\delta$ | 0.577 | 1.067 | 0.307 | 0.315 | 0.449 | 0.434 |
| $\delta$ | 0.705 | 1.858 | 0.088 | 0.436 | 1.056 | 0.180 |
| $\delta$ | 0.833 | -0.356 | 0.728 | 0.120 | 0.294 | 0.809 |
| $\delta$ | 0.962 | 3.250 | 0.007 | 0.566 | 7.793 | 0.033 |

Continued on next page



This work is shared under a CC BY license unless otherwise noted

Table 4: **Wave direction statistics**

| Mode | Angle | $T$ | $p$ | $d$ | $BF_{10}$ | $p_{FDR}$ |
|---|---|---|---|---|---|---|
| $\delta$ | 1.090 | 1.189 | 0.258 | 0.219 | 0.500 | 0.379 |
| $\delta$ | 1.218 | 0.163 | 0.873 | 0.038 | 0.282 | 0.897 |
| $\delta$ | 1.346 | -1.996 | 0.069 | 0.618 | 1.262 | 0.153 |
| $\delta$ | 1.475 | -2.824 | 0.015 | 1.067 | 4.092 | 0.055 |
| $\delta$ | 1.603 | -2.224 | 0.046 | 0.943 | 1.718 | 0.113 |
| $\delta$ | 1.731 | -1.785 | 0.099 | 0.668 | 0.964 | 0.194 |
| $\delta$ | 1.859 | -0.249 | 0.808 | 0.085 | 0.286 | 0.861 |
| $\delta$ | 1.988 | 1.877 | 0.085 | 0.677 | 1.082 | 0.177 |
| $\delta$ | 2.116 | 2.655 | 0.021 | 1.172 | 3.185 | 0.066 |
| $\delta$ | 2.244 | 2.165 | 0.051 | 0.947 | 1.583 | 0.122 |
| $\delta$ | 2.372 | -0.854 | 0.410 | 0.375 | 0.380 | 0.544 |
| $\delta$ | 2.500 | 0.558 | 0.587 | 0.228 | 0.319 | 0.709 |
| $\delta$ | 2.629 | 0.256 | 0.802 | 0.122 | 0.286 | 0.861 |
| $\delta$ | 2.757 | 0.399 | 0.697 | 0.183 | 0.298 | 0.780 |
| $\delta$ | 2.885 | 0.246 | 0.810 | 0.110 | 0.286 | 0.861 |
| $\delta$ | 3.013 | -0.182 | 0.859 | 0.077 | 0.282 | 0.888 |
| $\delta$ | 3.142 | -2.418 | 0.032 | 0.821 | 2.259 | 0.092 |
| S | -3.142 | -5.027 | 2.96e-04 | 1.987 | 1.13e+02 | 0.007 |
| S | -3.013 | 0.143 | 0.888 | 0.066 | 0.281 | 0.905 |
| S | -2.885 | 1.104 | 0.291 | 0.485 | 0.463 | 0.416 |
| S | -2.757 | 1.147 | 0.274 | 0.493 | 0.482 | 0.395 |
| S | -2.629 | 1.676 | 0.120 | 0.730 | 0.844 | 0.220 |
| S | -2.500 | 0.643 | 0.532 | 0.238 | 0.333 | 0.667 |
| S | -2.372 | -1.341 | 0.205 | 0.534 | 0.581 | 0.319 |
| S | -2.244 | 3.013 | 0.011 | 1.001 | 5.437 | 0.044 |
| S | -2.116 | 2.078 | 0.060 | 0.714 | 1.407 | 0.136 |
| S | -1.988 | 1.348 | 0.203 | 0.414 | 0.585 | 0.319 |
| S | -1.859 | 0.567 | 0.581 | 0.163 | 0.320 | 0.709 |
| S | -1.731 | -2.817 | 0.016 | 1.106 | 4.053 | 0.055 |
| S | -1.603 | -4.275 | 0.001 | 1.748 | 37.141 | 0.012 |
| S | -1.475 | -1.812 | 0.095 | 0.729 | 0.997 | 0.190 |
| S | -1.346 | -0.399 | 0.697 | 0.195 | 0.298 | 0.780 |
| S | -1.218 | 1.703 | 0.114 | 0.660 | 0.872 | 0.214 |
| S | -1.090 | 1.635 | 0.128 | 0.646 | 0.804 | 0.226 |
| S | -0.962 | 1.664 | 0.122 | 0.713 | 0.832 | 0.220 |
| S | -0.833 | -1.584 | 0.139 | 0.640 | 0.757 | 0.240 |
| S | -0.705 | -0.206 | 0.840 | 0.084 | 0.283 | 0.882 |
| S | -0.577 | 1.904 | 0.081 | 0.550 | 1.119 | 0.172 |
| S | -0.449 | 2.682 | 0.020 | 0.648 | 3.315 | 0.066 |
| S | -0.321 | 4.165 | 0.001 | 1.081 | 31.490 | 0.012 |
| S | -0.192 | 0.432 | 0.673 | 0.173 | 0.302 | 0.776 |
| S | -0.064 | -3.210 | 0.007 | 1.324 | 7.332 | 0.034 |

Continued on next page



This work is shared under a CC BY license unless otherwise noted

Table 4: **Wave direction statistics**

| Mode | Angle | $T$ | $p$ | $d$ | $BF_{10}$ | $p_{FDR}$ |
|---|---|---|---|---|---|---|
| S | 0.064 | -2.662 | 0.021 | 1.128 | 3.217 | 0.066 |
| S | 0.192 | 0.548 | 0.594 | 0.224 | 0.317 | 0.709 |
| S | 0.321 | 1.411 | 0.184 | 0.589 | 0.625 | 0.297 |
| S | 0.449 | 4.351 | 9.44e-04 | 1.318 | 41.624 | 0.012 |
| S | 0.577 | 4.191 | 0.001 | 1.521 | 32.752 | 0.012 |
| S | 0.705 | 3.545 | 0.004 | 1.489 | 12.243 | 0.025 |
| S | 0.833 | 0.481 | 0.639 | 0.209 | 0.308 | 0.743 |
| S | 0.962 | 5.808 | 8.37e-05 | 1.695 | 3.37e+02 | 0.003 |
| S | 1.090 | 3.861 | 0.002 | 1.457 | 19.852 | 0.016 |
| S | 1.218 | 1.739 | 0.108 | 0.650 | 0.910 | 0.207 |
| S | 1.346 | -0.406 | 0.692 | 0.175 | 0.299 | 0.780 |
| S | 1.475 | -2.913 | 0.013 | 1.126 | 4.682 | 0.050 |
| S | 1.603 | -3.988 | 0.002 | 1.509 | 24.084 | 0.015 |
| S | 1.731 | -1.921 | 0.079 | 0.795 | 1.144 | 0.169 |
| S | 1.859 | 0.510 | 0.619 | 0.209 | 0.312 | 0.726 |
| S | 1.988 | 1.540 | 0.149 | 0.747 | 0.720 | 0.255 |
| S | 2.116 | 2.042 | 0.064 | 0.971 | 1.342 | 0.143 |
| S | 2.244 | 1.336 | 0.206 | 0.595 | 0.578 | 0.319 |
| S | 2.372 | -4.190 | 0.001 | 1.073 | 32.680 | 0.012 |
| S | 2.500 | -0.793 | 0.443 | 0.271 | 0.364 | 0.577 |
| S | 2.629 | 1.147 | 0.274 | 0.489 | 0.482 | 0.395 |
| S | 2.757 | 0.787 | 0.446 | 0.339 | 0.363 | 0.577 |
| S | 2.885 | 0.759 | 0.462 | 0.308 | 0.356 | 0.593 |
| S | 3.013 | -0.324 | 0.752 | 0.120 | 0.291 | 0.823 |
| S | 3.142 | -5.148 | 2.42e-04 | 1.415 | 1.34e+02 | 0.007 |
| U-S | -3.142 | -2.741 | 0.018 | 1.010 | 3.617 | 0.062 |
| U-S | -3.013 | 2.557 | 0.025 | 1.034 | 2.758 | 0.075 |
| U-S | -2.885 | 3.266 | 0.007 | 1.222 | 7.992 | 0.033 |
| U-S | -2.757 | 3.076 | 0.010 | 1.199 | 5.989 | 0.041 |
| U-S | -2.629 | 3.245 | 0.007 | 1.150 | 7.742 | 0.033 |
| U-S | -2.500 | 1.046 | 0.316 | 0.388 | 0.441 | 0.439 |
| U-S | -2.372 | -2.442 | 0.031 | 0.936 | 2.337 | 0.090 |
| U-S | -2.244 | 4.389 | 8.82e-04 | 1.728 | 44.086 | 0.012 |
| U-S | -2.116 | 2.241 | 0.045 | 0.826 | 1.760 | 0.113 |
| U-S | -1.988 | 1.215 | 0.248 | 0.445 | 0.513 | 0.372 |
| U-S | -1.859 | -0.539 | 0.599 | 0.221 | 0.316 | 0.709 |
| U-S | -1.731 | -4.478 | 7.56e-04 | 1.746 | 50.302 | 0.012 |
| U-S | -1.603 | -10.375 | 2.40e-07 | 2.639 | 6.11e+04 | 3.61e-05 |
| U-S | -1.475 | -7.412 | 8.15e-06 | 2.453 | 2.62e+03 | 4.07e-04 |
| U-S | -1.346 | -1.661 | 0.123 | 0.564 | 0.829 | 0.220 |
| U-S | -1.218 | 1.021 | 0.328 | 0.348 | 0.432 | 0.451 |
| U-S | -1.090 | 3.567 | 0.004 | 1.069 | 12.662 | 0.025 |

Continued on next page



This work is shared under a CC BY license unless otherwise noted

Table 4: **Wave direction statistics**

| Mode | Angle | $T$ | $p$ | $d$ | $BF_{10}$ | $p_{FDR}$ |
|---|---|---|---|---|---|---|
| U-S | -0.962 | 4.726 | 4.92e-04 | 1.452 | 72.641 | 0.010 |
| U-S | -0.833 | -1.803 | 0.097 | 0.709 | 0.985 | 0.191 |
| U-S | -0.705 | 0.538 | 0.600 | 0.213 | 0.316 | 0.709 |
| U-S | -0.577 | 4.435 | 8.14e-04 | 1.893 | 47.202 | 0.012 |
| U-S | -0.449 | 3.942 | 0.002 | 1.751 | 22.452 | 0.015 |
| U-S | -0.321 | 2.826 | 0.015 | 1.195 | 4.104 | 0.055 |
| U-S | -0.192 | 2.547 | 0.026 | 0.993 | 2.720 | 0.075 |
| U-S | -0.064 | -3.883 | 0.002 | 0.772 | 20.532 | 0.016 |
| U-S | 0.064 | -3.841 | 0.002 | 0.835 | 19.250 | 0.016 |
| U-S | 0.192 | 1.717 | 0.112 | 0.607 | 0.886 | 0.212 |
| U-S | 0.321 | 0.928 | 0.372 | 0.333 | 0.401 | 0.498 |
| U-S | 0.449 | 2.117 | 0.056 | 0.719 | 1.484 | 0.131 |
| U-S | 0.577 | 2.172 | 0.051 | 0.669 | 1.600 | 0.122 |
| U-S | 0.705 | 0.641 | 0.534 | 0.238 | 0.332 | 0.667 |
| U-S | 0.833 | -1.659 | 0.123 | 0.798 | 0.827 | 0.220 |
| U-S | 0.962 | 2.309 | 0.040 | 0.836 | 1.935 | 0.106 |
| U-S | 1.090 | 1.354 | 0.201 | 0.522 | 0.589 | 0.319 |
| U-S | 1.218 | 0.561 | 0.585 | 0.233 | 0.319 | 0.709 |
| U-S | 1.346 | -0.566 | 0.582 | 0.254 | 0.320 | 0.709 |
| U-S | 1.475 | -4.665 | 5.46e-04 | 1.881 | 66.439 | 0.010 |
| U-S | 1.603 | -9.065 | 1.02e-06 | 2.648 | 1.66e+04 | 7.67e-05 |
| U-S | 1.731 | -3.447 | 0.005 | 1.274 | 10.536 | 0.029 |
| U-S | 1.859 | 0.190 | 0.852 | 0.071 | 0.283 | 0.888 |
| U-S | 1.988 | 0.800 | 0.439 | 0.319 | 0.366 | 0.577 |
| U-S | 2.116 | 1.282 | 0.224 | 0.570 | 0.548 | 0.339 |
| U-S | 2.244 | 1.827 | 0.093 | 0.793 | 1.015 | 0.188 |
| U-S | 2.372 | -2.639 | 0.022 | 1.195 | 3.110 | 0.066 |
| U-S | 2.500 | 0.970 | 0.351 | 0.445 | 0.414 | 0.475 |
| U-S | 2.629 | 3.059 | 0.010 | 1.227 | 5.831 | 0.041 |
| U-S | 2.757 | 3.068 | 0.010 | 1.222 | 5.912 | 0.041 |
| U-S | 2.885 | 3.001 | 0.011 | 1.094 | 5.342 | 0.044 |
| U-S | 3.013 | 3.270 | 0.007 | 1.192 | 8.043 | 0.033 |
| U-S | 3.142 | -4.270 | 0.001 | 1.362 | 36.870 | 0.012 |

Table 5: **Recurrence network topology statistics**

| Measure | Mode | $T$ | $p$ | $d$ | $BF_{10}$ | $p_{FDR}$ |
|---|---|---|---|---|---|---|
| ge | $\delta$ | -4.402 | 8.62e-04 | 1.769 | 44.936 | 0.001 |
| $N_{coms}$ | $\delta$ | 6.288 | 4.02e-05 | 2.647 | 6.40e+02 | 6.89e-05 |
| $Q$ | $\delta$ | 3.806 | 0.003 | 1.516 | 18.234 | 0.003 |
| $NE$ | $\delta$ | 6.051 | 5.75e-05 | 2.516 | 4.67e+02 | 8.63e-05 |
| ge | S | -6.698 | 2.20e-05 | 2.487 | 1.09e+03 | 6.61e-05 |

Continued on next page



This work is shared under a CC BY license unless otherwise noted

Table 5: **Recurrence network topology statistics**

| Measure | Mode | $T$ | $p$ | $d$ | $BF_{10}$ | $p_{FDR}$ |
|---|---|---|---|---|---|---|
| $N_{coms}$ | S | 3.902 | 0.002 | 1.498 | 21.129 | 0.003 |
| $Q$ | S | 6.439 | 3.21e-05 | 2.374 | 7.80e+02 | 6.89e-05 |
| $NE$ | S | 6.327 | 3.79e-05 | 2.322 | 6.73e+02 | 6.89e-05 |
| ge | U-S | -7.977 | 3.87e-06 | 2.206 | 5.07e+03 | 2.32e-05 |
| $N_{coms}$ | U-S | 2.416 | 0.033 | 0.792 | 2.252 | 0.033 |
| $Q$ | U-S | 9.612 | 5.48e-07 | 2.426 | 2.91e+04 | 6.58e-06 |
| $NE$ | U-S | 7.546 | 6.81e-06 | 2.287 | 3.07e+03 | 2.72e-05 |

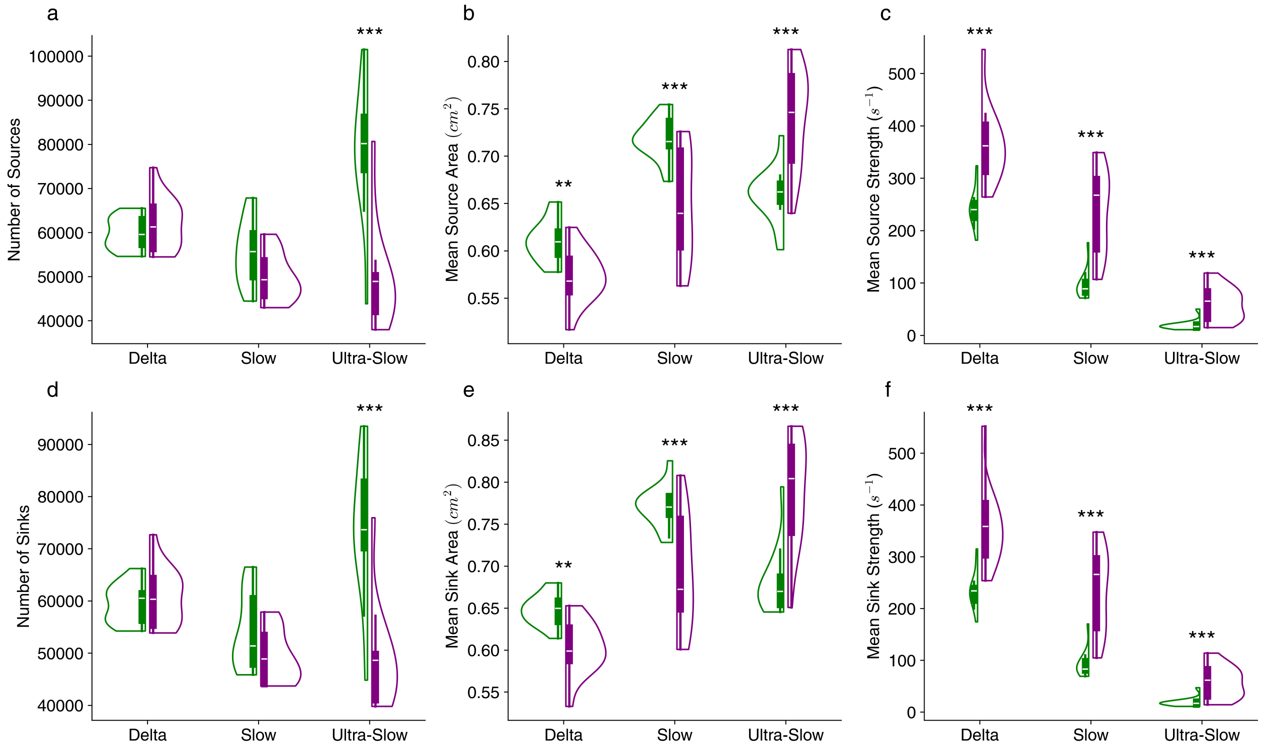


Figure 4: **Statistical summaries of flow field singularities** (*,**,*** indicates p<.05,.01,.001 FDR-corrected Benjamini–Hochberg procedure)

Table 6: **Singularity statistics**

| Measure | Mode | $T$ | $p$ | $d$ | $BF_{10}$ | $p_{FDR}$ |
|---|---|---|---|---|---|---|
| $N_{sources}$ | $\delta$ | 1.477 | 0.165 | 0.451 | 0.671 | 0.186 |
| $N_{sinks}$ | $\delta$ | 0.914 | 0.379 | 0.276 | 0.397 | 0.389 |
| $A(source)$ | $\delta$ | -3.355 | 0.006 | 1.497 | 9.150 | 0.008 |
| $A(sink)$ | $\delta$ | -4.381 | 8.95e-04 | 1.778 | 43.544 | 0.001 |
| $\nabla \cdot \vec{\mathbf{v}}(source)$ | $\delta$ | 4.939 | 3.42e-04 | 2.113 | 99.180 | 7.71e-04 |
| $\nabla \cdot \vec{\mathbf{v}}(sink)$ | $\delta$ | 4.939 | 3.43e-04 | 2.090 | 99.123 | 7.71e-04 |

Continued on next page



This work is shared under a CC BY license unless otherwise noted

Table 6: **Singularity statistics**

| Measure | Mode | $T$ | $p$ | $d$ | $BF_{10}$ | $p_{FDR}$ |
|---|---|---|---|---|---|---|
| $\nu$ | $\delta$ | 5.503 | 1.36e-04 | 2.279 | 2.21e+02 | 5.23e-04 |
| $A(asym)$ | $\delta$ | 2.393 | 0.034 | 0.668 | 2.180 | 0.046 |
| $A\nabla \cdot \vec{\mathbf{v}}(asym)$ | $\delta$ | 1.074 | 0.304 | 0.403 | 0.452 | 0.328 |
| $N_{sources}$ | S | -2.289 | 0.041 | 0.861 | 1.881 | 0.053 |
| $N_{sinks}$ | S | -2.138 | 0.054 | 0.805 | 1.527 | 0.063 |
| $A(source)$ | S | -4.682 | 5.30e-04 | 1.578 | 68.116 | 9.54e-04 |
| $A(sink)$ | S | -4.733 | 4.86e-04 | 1.536 | 73.445 | 9.37e-04 |
| $\nabla \cdot \vec{\mathbf{v}}(source)$ | S | 6.239 | 4.32e-05 | 2.308 | 6.00e+02 | 2.34e-04 |
| $\nabla \cdot \vec{\mathbf{v}}(sink)$ | S | 6.270 | 4.13e-05 | 2.317 | 6.25e+02 | 2.34e-04 |
| $\nu$ | S | 6.549 | 2.73e-05 | 2.397 | 8.98e+02 | 2.34e-04 |
| $A(asym)$ | S | 0.894 | 0.389 | 0.367 | 0.391 | 0.389 |
| $A\nabla \cdot \vec{\mathbf{v}}(asym)$ | S | 3.457 | 0.005 | 1.375 | 10.699 | 0.007 |
| $N_{sources}$ | U-S | -6.454 | 3.14e-05 | 2.213 | 7.95e+02 | 2.34e-04 |
| $N_{sinks}$ | U-S | -6.277 | 4.08e-05 | 2.155 | 6.31e+02 | 2.34e-04 |
| $A(source)$ | U-S | 4.829 | 4.13e-04 | 1.861 | 84.494 | 8.57e-04 |
| $A(sink)$ | U-S | 5.506 | 1.35e-04 | 1.926 | 2.22e+02 | 5.23e-04 |
| $\nabla \cdot \vec{\mathbf{v}}(source)$ | U-S | 5.076 | 2.72e-04 | 1.675 | 1.21e+02 | 7.35e-04 |
| $\nabla \cdot \vec{\mathbf{v}}(sink)$ | U-S | 5.086 | 2.68e-04 | 1.689 | 1.23e+02 | 7.35e-04 |
| $\nu$ | U-S | 5.176 | 2.31e-04 | 1.684 | 1.39e+02 | 7.35e-04 |
| $A(asym)$ | U-S | -2.235 | 0.045 | 0.931 | 1.744 | 0.056 |
| $A\nabla \cdot \vec{\mathbf{v}}(asym)$ | U-S | 4.387 | 8.85e-04 | 1.706 | 43.962 | 0.001 |

Table 7: **Wave pattern statistics**

| Measure | Mode | $\text{Wave}_{Type}$ | $T$ | $p$ | $d$ | $BF_{10}$ | $p_{FDR}$ |
|---|---|---|---|---|---|---|---|
| Lifetime | $\delta$ | S-Node | -5.350 | 1.74e-04 | 2.261 | 1.78e+02 | 6.01e-04 |
| Lifetime | $\delta$ | S-Focus | -5.880 | 7.49e-05 | 2.210 | 3.71e+02 | 3.31e-04 |
| Lifetime | $\delta$ | U-Node | -4.547 | 6.70e-04 | 1.905 | 55.747 | 0.002 |
| Lifetime | $\delta$ | U-Focus | -5.179 | 2.30e-04 | 2.169 | 1.40e+02 | 7.38e-04 |
| Lifetime | $\delta$ | Saddle | -4.119 | 0.001 | 1.452 | 29.362 | 0.003 |
| Proportion | $\delta$ | S-Node | 2.085 | 0.059 | 0.676 | 1.421 | 0.081 |
| Proportion | $\delta$ | S-Focus | 0.768 | 0.457 | 0.231 | 0.358 | 0.485 |
| Proportion | $\delta$ | U-Node | 1.934 | 0.077 | 0.643 | 1.164 | 0.099 |
| Proportion | $\delta$ | U-Focus | 0.432 | 0.673 | 0.137 | 0.302 | 0.673 |
| Proportion | $\delta$ | Saddle | 1.357 | 0.200 | 0.405 | 0.590 | 0.219 |
| Number | $\delta$ | S-Node | 3.776 | 0.003 | 1.331 | 17.426 | 0.005 |
| Number | $\delta$ | S-Focus | 2.324 | 0.038 | 0.697 | 1.976 | 0.062 |
| Number | $\delta$ | U-Node | 4.300 | 0.001 | 1.392 | 38.581 | 0.002 |
| Number | $\delta$ | U-Focus | 2.219 | 0.047 | 0.710 | 1.705 | 0.070 |
| Number | $\delta$ | Saddle | 1.853 | 0.089 | 0.561 | 1.049 | 0.111 |
| Lifetime | S | S-Node | -8.757 | 1.47e-06 | 2.744 | 1.20e+04 | 6.63e-05 |
| Lifetime | S | S-Focus | -6.183 | 4.70e-05 | 2.148 | 5.57e+02 | 2.65e-04 |

Continued on next page



This work is shared under a CC BY license unless otherwise noted

Table 7: **Wave pattern statistics**

| Measure | Mode | $Wave_{Type}$ | $T$ | $p$ | $d$ | $BF_{10}$ | $p_{FDR}$ |
|---|---|---|---|---|---|---|---|
| Lifetime | S | U-Node | -6.950 | 1.54e-05 | 2.601 | 1.49e+03 | 1.73e-04 |
| Lifetime | S | U-Focus | -7.246 | 1.02e-05 | 2.239 | 2.15e+03 | 1.73e-04 |
| Lifetime | S | Saddle | -1.524 | 0.153 | 0.486 | 0.707 | 0.173 |
| Proportion | S | S-Node | -1.943 | 0.076 | 0.567 | 1.177 | 0.099 |
| Proportion | S | S-Focus | -2.761 | 0.017 | 0.940 | 3.725 | 0.031 |
| Proportion | S | U-Node | -1.571 | 0.142 | 0.642 | 0.746 | 0.164 |
| Proportion | S | U-Focus | -2.195 | 0.049 | 0.870 | 1.650 | 0.070 |
| Proportion | S | Saddle | -2.087 | 0.059 | 0.812 | 1.426 | 0.081 |
| Number | S | S-Node | 3.649 | 0.003 | 0.951 | 14.355 | 0.007 |
| Number | S | S-Focus | -0.758 | 0.463 | 0.245 | 0.356 | 0.485 |
| Number | S | U-Node | 2.227 | 0.046 | 0.842 | 1.725 | 0.070 |
| Number | S | U-Focus | -0.499 | 0.627 | 0.182 | 0.310 | 0.641 |
| Number | S | Saddle | -1.680 | 0.119 | 0.641 | 0.847 | 0.145 |
| Lifetime | U-S | S-Node | -4.619 | 5.91e-04 | 1.482 | 62.056 | 0.002 |
| Lifetime | U-S | S-Focus | 3.522 | 0.004 | 1.107 | 11.818 | 0.008 |
| Lifetime | U-S | U-Node | -2.450 | 0.031 | 1.039 | 2.362 | 0.051 |
| Lifetime | U-S | U-Focus | 1.653 | 0.124 | 0.588 | 0.821 | 0.147 |
| Lifetime | U-S | Saddle | 5.463 | 1.45e-04 | 2.256 | 2.09e+02 | 5.42e-04 |
| Proportion | U-S | S-Node | -5.829 | 8.10e-05 | 1.778 | 3.47e+02 | 3.31e-04 |
| Proportion | U-S | S-Focus | -6.968 | 1.50e-05 | 2.189 | 1.52e+03 | 1.73e-04 |
| Proportion | U-S | U-Node | -5.952 | 6.70e-05 | 2.235 | 4.09e+02 | 3.31e-04 |
| Proportion | U-S | U-Focus | -5.105 | 2.60e-04 | 1.737 | 1.26e+02 | 7.79e-04 |
| Proportion | U-S | Saddle | -6.429 | 3.26e-05 | 2.321 | 7.69e+02 | 2.29e-04 |
| Number | U-S | S-Node | -2.632 | 0.022 | 0.945 | 3.080 | 0.038 |
| Number | U-S | S-Focus | -6.590 | 2.58e-05 | 1.989 | 9.47e+02 | 2.29e-04 |
| Number | U-S | U-Node | -4.273 | 0.001 | 1.657 | 37.019 | 0.002 |
| Number | U-S | U-Focus | -4.636 | 5.75e-04 | 1.529 | 63.591 | 0.002 |
| Number | U-S | Saddle | -6.370 | 3.56e-05 | 2.319 | 7.13e+02 | 2.29e-04 |



This work is shared under a CC BY license unless otherwise noted

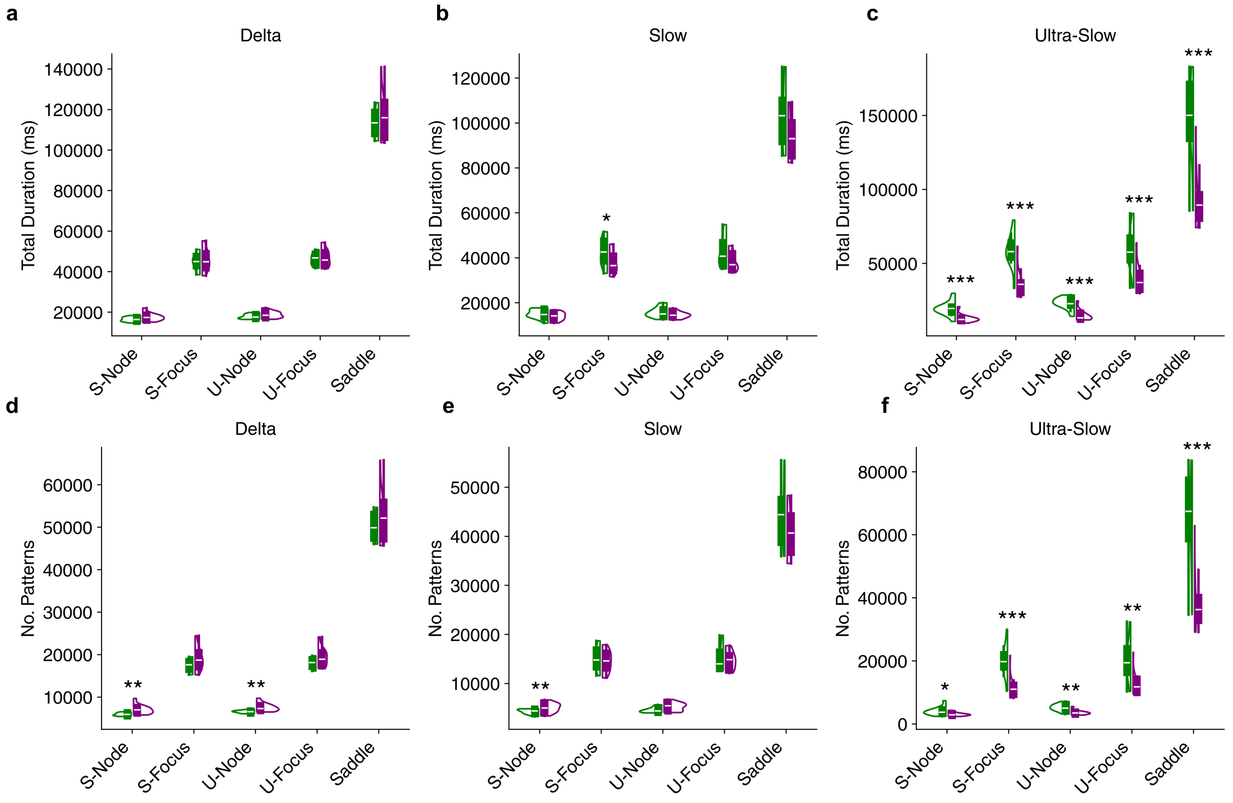


Figure 5: **Extended pattern temporal dynamics a,b,c** Total life times of waves **d,e,f** Number of unique waves observed. (*,**,*** indicates p<.05,.01,.001 FDR-corrected Benjamini–Hochberg procedure)

Table 8: **Timescale statistics**

| Measure | T | $p$ | $d$ | $BF_{10}$ | $p_{FDR}$ |
|---|---|---|---|---|---|
| $\lambda_{max}$ | -2.221 | 0.046 | 0.724 | 1.712 | 0.046 |
| $IT$ | 3.125 | 0.009 | 0.898 | 6.446 | 0.018 |

Table 9: **SVD-PCA statistics**

| Measure | T | $p$ | $d$ | $BF_{10}$ | $p_{FDR}$ |
|---|---|---|---|---|---|
| $Eigenvector1$ | 2.514 | 0.027 | 0.911 | 2.592 | 0.027 |
| $Eigenvector2$ | 3.640 | 0.003 | 0.899 | 14.162 | 0.004 |
| $Eigenvector3$ | 3.508 | 0.004 | 1.054 | 11.576 | 0.005 |
| $Eigenvector4$ | 3.941 | 0.002 | 1.151 | 22.426 | 0.004 |
| $Eigenvector5$ | 3.691 | 0.003 | 1.081 | 15.301 | 0.004 |
| $Eigenvector6$ | 3.628 | 0.003 | 1.202 | 13.899 | 0.004 |
| $Eigenvector7$ | 3.922 | 0.002 | 1.296 | 21.760 | 0.004 |
| $Eigenvector8$ | 3.808 | 0.002 | 1.301 | 18.304 | 0.004 |

Continued on next page



This work is shared under a CC BY license unless otherwise noted

Table 9: **SVD-PCA statistics**

| Measure | T | $p$ | $d$ | $BF_{10}$ | $p_{FDR}$ |
|---|---|---|---|---|---|
| $Eigenvector9$ | 3.955 | 0.002 | 1.354 | 22.900 | 0.004 |
| $Eigenvector10$ | 4.045 | 0.002 | 1.380 | 26.251 | 0.004 |

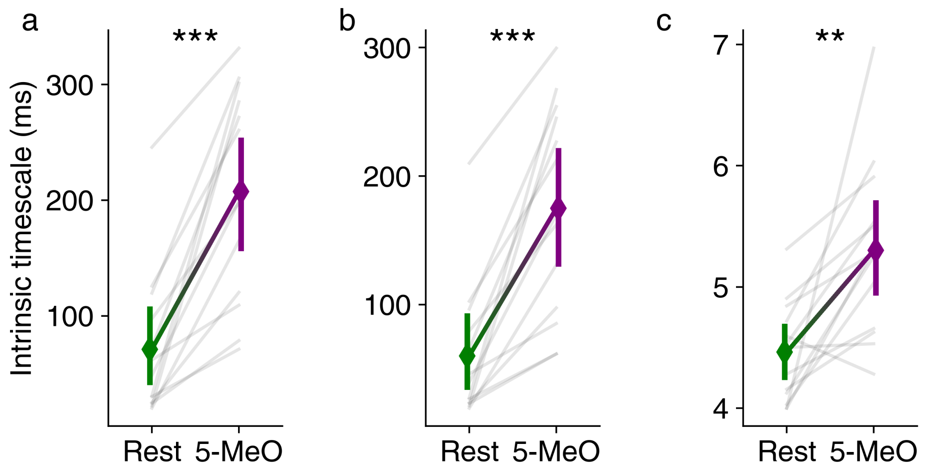


Figure 6: **Robustness of intrinsic timescales to parameter choices** **a** Decay point as 2 S.D above the stable minima **b** Decay point as 3 S.D above the stable minima **c** Decay point as 50% of original $AMI$. (*,**,*** indicates p<.05,.01,.001 FDR-corrected Benjamini–Hochberg procedure)

Table 10: **Energy landscape statistics**

| Msd | $T$ | $p$ | $BF_{10}$ | $d$ | $p_{FDR}$ |
|---|---|---|---|---|---|
| 1.000 | -4.628 | 5.82e-04 | 62.859 | 1.307 | 0.002 |
| 1.200 | -3.510 | 0.004 | 11.603 | 1.131 | 0.008 |
| 1.400 | -1.660 | 0.123 | 0.827 | 0.648 | 0.143 |
| 1.600 | -0.172 | 0.866 | 0.282 | 0.072 | 0.866 |
| 1.800 | 0.803 | 0.438 | 0.367 | 0.327 | 0.460 |
| 2.000 | 1.460 | 0.170 | 0.659 | 0.566 | 0.188 |
| 2.200 | 1.940 | 0.076 | 1.173 | 0.715 | 0.094 |
| 2.400 | 2.320 | 0.039 | 1.964 | 0.814 | 0.051 |
| 2.600 | 2.641 | 0.022 | 3.121 | 0.885 | 0.030 |
| 2.800 | 2.928 | 0.013 | 4.789 | 0.937 | 0.019 |
| 3.000 | 3.198 | 0.008 | 7.200 | 0.978 | 0.012 |
| 3.200 | 3.460 | 0.005 | 10.750 | 1.010 | 0.008 |
| 3.400 | 3.725 | 0.003 | 16.125 | 1.038 | 0.006 |
| 3.600 | 4.001 | 0.002 | 24.559 | 1.063 | 0.004 |
| 3.800 | 4.294 | 0.001 | 38.248 | 1.087 | 0.003 |
| 4.000 | 4.605 | 6.06e-04 | 60.760 | 1.111 | 0.002 |
| 4.200 | 4.919 | 3.54e-04 | 96.357 | 1.135 | 0.001 |

Continued on next page



This work is shared under a CC BY license unless otherwise noted

Table 10: **Energy landscape statistics**

| Msd | $T$ | $p$ | $BF_{10}$ | $d$ | $p_{FDR}$ |
|---|---|---|---|---|---|
| 4.400 | 5.207 | 2.19e-04 | 1.46e+02 | 1.158 | 0.001 |
| 4.600 | 5.423 | 1.54e-04 | 1.98e+02 | 1.178 | 0.001 |
| 4.800 | 5.527 | 1.30e-04 | 2.29e+02 | 1.193 | 0.001 |
| 5.000 | 5.497 | 1.37e-04 | 2.19e+02 | 1.200 | 0.001 |



This work is shared under a CC BY license unless otherwise noted
